# FOS binding sites are a hub for the evolution of activity-dependent gene regulatory programs in human neurons

**DOI:** 10.1101/2025.03.31.646366

**Authors:** Ava C. Carter, Gabriel T. Koreman, Jillian E. Petrocelli, Josephine E. Robb, Evan M. Bushinsky, Sara K. Trowbridge, David M. Kingsley, Christopher A. Walsh, Janet H.T. Song, Michael E. Greenberg

## Abstract

After birth, sensory inputs to neurons trigger the induction of activity-dependent genes (ADGs) that mediate many aspects of neuronal maturation and plasticity. To identify human-specific ADGs, we characterized these genes in human-chimpanzee tetraploid neurons. We identified 235 ADGs that are differentially expressed between human and chimpanzee neurons and found that their nearby regulatory sites are species-biased in their binding of the transcription factor FOS. An assessment of these sites revealed that many are enriched for single nucleotide variants that promote or eliminate FOS binding in human neurons. Disrupting the function of individual species-biased FOS-bound enhancers diminishes expression of nearby genes and affects the firing dynamics of human neurons. Our findings indicate that FOS-bound enhancers are frequent sites of evolution and that they regulate human-specific ADGs that may contribute to the unusually protracted and complex process of postnatal human brain development.

## Introduction

Compared to other species, the human brain displays differences in size, connectivity, and morphology that are thought to underlie our unique cognitive capabilities. One striking feature of human brain development that may contribute to these differences is the slow pace of neuronal maturation that continues into the third decade of life. Human cortical neurons show the largest degree of neoteny, the retention of juvenile features, and mature more slowly than those of their closest living relatives, the chimpanzees (Vanderhaeghen and Polleux, 2023). Notably, this slowed pace of maturation is preserved when human cortical neurons are transplanted into rodent brains, demonstrating that this neoteny is intrinsic to the neuron and genetically encoded (Gaspard et al., 2008; Espuny-Camacho et al., 2013; Otani et al., 2016).

The process of brain maturation and plasticity has been best studied in the mouse, where sensory-driven neuronal activity induces activity-dependent gene expression (ADGE) via calcium influx through L-type voltage sensitive calcium channels. ADGE regulates many aspects of brain maturation, including synaptic pruning, dendritic outgrowth, and excitatory-inhibitory balance, as well as the brain plasticity that underlies learning, memory, and behavior (Lin et al., 2008; Gu et al., 2012; Favuzzi et al., 2017; Yap and Greenberg, 2018; Yap et al., 2021). There are two waves of ADGE. In the first wave, immediate early genes (IEGs), which encode transcription factors (TFs), including the AP-1 factor FOS, are rapidly and transiently induced. These TFs promote the induction of a second wave of late response genes (LRGs), many of which encode secreted proteins that regulate critical aspects of brain maturation (Holliday et al., 1991; Carey and Matsumoto, 1999; Deisseroth et al., 2004; Ringler et al., 2008; Piatti et al., 2011; Faria-Pereira et al., 2022; Hergenreder et al., 2024).

Given the importance of neuronal activity for brain plasticity and the dramatically protracted pace of neuronal maturation in humans, we hypothesized that the ADGE program evolved unique features in humans. Prior studies identified genes, including *Osteocrin*, that are activity-dependent in human and primate neurons, but not rodent neurons, and suggested that these genes may have become activity-dependent through evolution by the gain of nearby activity-dependent regulatory regions (Ataman et al., 2016; Qiu et al., 2016; Pruunsild et al., 2017). Activity-dependent regulatory regions have also been shown to have heritability enrichment for neurological traits including autism spectrum disorder, schizophrenia, and general intelligence, underscoring the importance of identifying human-specific features of ADGE (Boulting et al., 2021; Sanchez-Priego et al., 2022).

Despite this progress, virtually nothing is known about the human-specific changes to ADGE that might account for the prolonged maturation and increased complexity of the human brain. The identification of such changes might be accomplished by a careful comparison of ADGE in human neurons and neurons from our closest living relative, the chimpanzee. The majority of the 20 million single nucleotide variants (SNVs) between the human and chimpanzee genomes occur in non-coding regions, suggesting that changes in the spatiotemporal regulation of gene expression account for many of the evolutionary changes between these two species (King and Wilson, 1975; Carroll, 2008; McLean et al., 2011). However, prioritizing single sequence variants and the specific gene regulatory regions that contribute to evolutionary changes in ADGE has not yet been achieved.

Recent advances in single-cell genomics have facilitated the identification of many gene expression differences between primates, including humans, within specific cell types of the brain (Jorstad et al., 2023; Caglayan et al., 2023). However, these pioneering studies have several limitations, including difficulty in accurately matching developmental stage and cell type across species, and the inability to stimulate neurons in postmortem tissue in a homogeneous and synchronized way. Thus, we aimed to study ADGE in a neuronal culture system in which we could directly compare gene transcription in human and chimpanzee neurons of the exact same developmental stage in response to a controlled, synchronized external stimulus.

To study differences in gene expression between human and chimpanzee neurons in the same nuclear environment, we used a human-chimpanzee tetraploid model (Agoglia et al., 2021; Gokhman et al., 2021; Song et al., 2021), which allowed us to distinguish between *cis*- and *trans*-regulatory effects. *Cis*-regulatory effects are driven by sequence differences between species in regulatory regions linked to a gene of interest, including promoters and enhancers. Characterizing these *cis*-regulated differences allowed us to test the effect of sequence changes on gene expression and ask whether they contribute to phenotypic differences between humans and chimpanzees. *Trans*-regulatory effects on gene expression, on the other hand, arise from species differences in the abundance of diffusible factors, such as the abundance of TFs that regulate promoters and enhancers, and frequently control large networks of unlinked target genes.

Here, we achieved the first detailed, mechanistic comparison of ADGE between humans and chimpanzees by differentiating human and chimpanzee diploid and tetraploid cells into excitatory neurons and applying a controlled, synchronized stimulus that activates ADGE (Figure 1a). We identified ADGs that are inducible in humans, but not chimpanzees, suggesting they have acquired activity-dependent regulation specifically in humans. Our analysis also revealed numerous genes that are inducible in both humans and chimpanzees but that show consistently higher expression in one of the two species across the time course. To decipher the evolutionary mechanisms that gave rise to the human-specific features of the ADGE program, we characterized the regulatory regions that control human-biased ADGE. We identified a preponderance of binding for the activity-dependent TF FOS at sites where SNVs between humans and chimpanzees appear to drive species-specific differences in the ADG response. We demonstrate that FOS binding site changes that occurred during the evolution of humans led to the creation of new activity-dependent enhancers that regulate genes such as *TUNAR*, whose expression affects the bursting dynamics of excitatory neurons.

**Figure 1:**
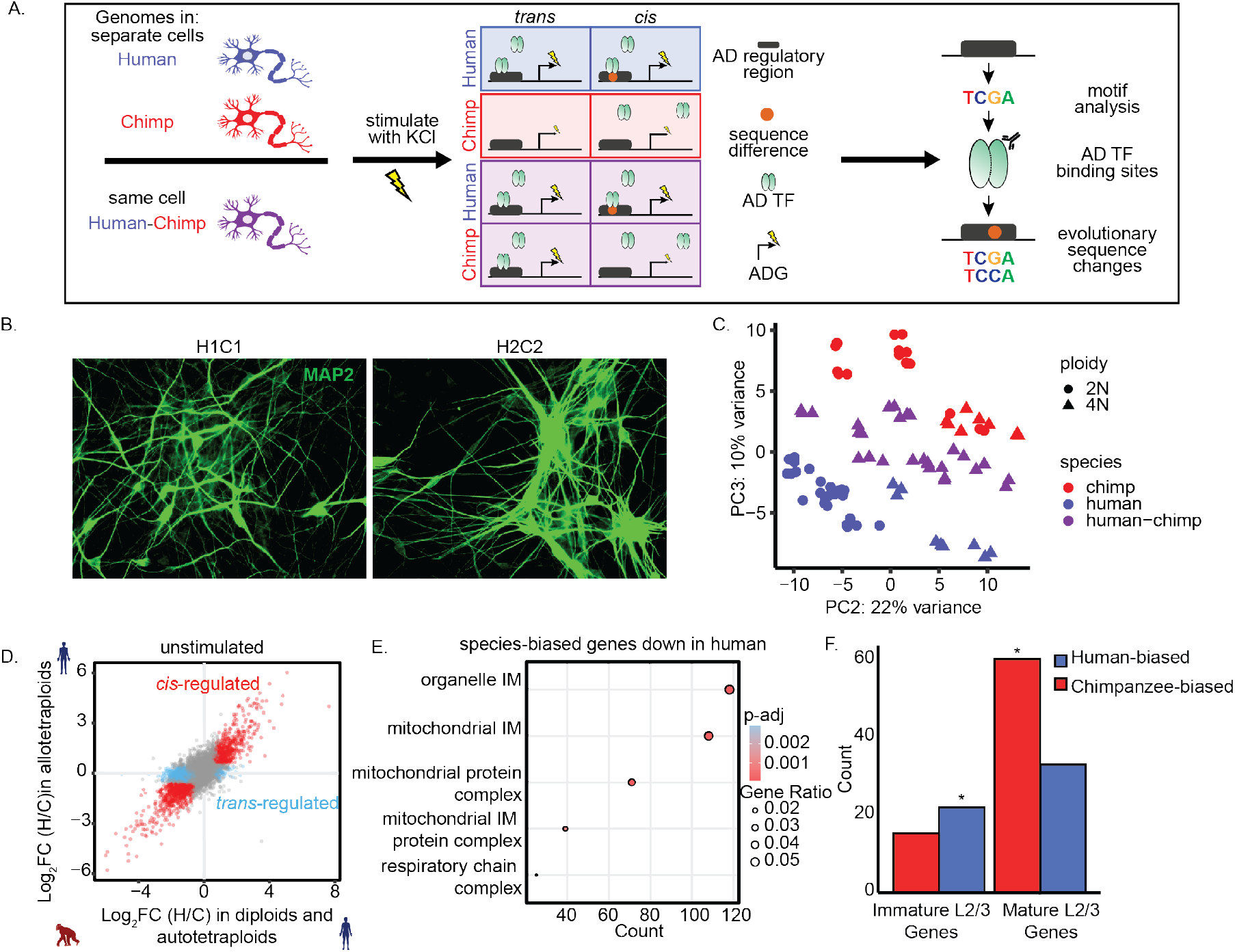
Identifying species-biased genes in human-chimpanzee tetraploid neurons. a. Diagram of human-chimpanzee tetraploid piNs and analysis pipeline. b. Immunofluorescence images of two human-chimpanzee allotetraploid iPSC lines differentiated into piNs for 28 days. Neurons are stained for MAP2 (green) to mark mature dendrites. c. PCA of RNA-seq data for diploid and tetraploid piNs showing PC2 vs. PC3. d. Scatterplot of RNA-seq data showing the log_2_ fold change in expression between human and chimpanzee (H/C) alleles in allotetraploid cells vs the log_2_ fold change in expression between human and chimpanzee neurons in diploid and autotetraploid cells. *Cis*-regulated genes are indicated in red and *trans*-regulated genes are indicated in blue. e. GO analysis for genes that are chimpanzee-biased in unstimulated piNs. IM= inner membrane. f. Enrichment of human-biased and chimp-biased genes in gene sets from Herring et al. 2022 that significantly decrease or increase in expression from fetal to neonatal timepoints in primary layer II/III CUX2+ neurons (* p = 0.0063; Hypergeometric test for overrepresentation).

## Results

### Human-chimpanzee tetraploid cells differentiate into excitatory neurons

To compare gene expression between human and chimpanzee alleles in cortical neurons, we generated human and chimpanzee diploid and tetraploid neurons by overexpressing the neurogenic TF NGN2 combined with dual SMAD inhibition in human or chimpanzee induced pluripotent stem cells (iPSCs) (Song et al., 2021). This protocol produces cells with a forebrain excitatory neuron-like phenotype that we refer to as patterned induced cortical neurons (piNs) (Nehme et al., 2018). The process of piN differentiation is highly reproducible and produces a relatively homogeneous population of neurons in 28 days. To generate these iPSC lines, we used CRISPR/Cas9 to insert a doxycycline-inducible expression cassette for NGN2 into the AAVS1 safe-harbor locus and selected cells with this insert for clonal expansion into stable cell lines. We generated multiple clones from each starting iPSC line (full cell line info in Table S1). The resulting piNs included diploids of each species, as well as human-human and chimpanzee-chimpanzee autotetraploids to control for tetraploidization effects, and human-chimpanzee allotetraploids to allow comparison of human and chimpanzee allelic gene regulation within the same cellular environment. At day 28 (D28) of differentiation, allotetraploid piNs derived from two iPSC lines expressed MAP2 and had clear neuronal morphology (Figure 1b), confirming that tetraploid iPSCs could differentiate into piNs that are similar in their phenotypic characteristics to diploid iPSC-derived piNs. Phenotypic variability across diploids, autotetraploids, and allotetraploids was not associated with differences in ploidy or species of origin (Figure S1).

To identify human-specific activity-dependent genes, we either left piNs untreated or exposed them to an elevated level of extracellular potassium chloride (KCl), which leads to membrane depolarization followed by the opening of L-type voltage-gated calcium channels and an influx of calcium that triggers the activation of the ADGE program (Sheng and Greenberg, 1990; Bading et al., 1993; Xia et al., 1996; Tao et al., 1998). It is noteworthy that, in mice, the KCl protocol is a well-established *in vitro* method for inducing an ADGE program that is similar to the ADGE that is activated *in vivo* in response to sensory stimuli (Lin et al., 2008; Hrvatin et al., 2018; Yap et al., 2021; Traunmüller et al., 2025).

Principal component (PC) analysis of the RNA-seq data obtained from uninduced and KCl treated piNs (human and chimpanzee diploids, autotetraploids, and allotetraploids) revealed that the samples separated in the first and second PCs by biological replicate (separate differentiations), and in the second and third PCs by species (Figure 1c, Figure S2a,b). Diploid and autotetraploid samples from the same species were partially separated in these PCs, indicating that expression of a subset of genes reflects the tetraploid state (see STAR Methods). All lines expressed cortical neuron markers at D28. Variability in the expression of previously described piN marker genes (Nehme et al., 2018) was not driven by ploidy, species, or experimental replicate (Figure S2c). Similarly, all lines expressed the IEG *FOS* after membrane depolarization, and any variability detected in *FOS* expression was not associated with ploidy, species, or experimental replicate (Figure S2d,e).

A major advantage of studying gene expression in the human-chimpanzee tetraploid piNs is the ability to separate *cis*-regulated changes in gene expression between human and chimpanzee that arise due to sequence changes within nearby regulatory elements from *trans*-regulated differences that reflect changes in the overall transcriptional environment. In this study, *cis*-regulated genes are identified as those that show significant species-specific expression differences when compared between separate human and chimpanzee diploid and autotetraploid piNs, and that still show similar allele-specific expression differences when the human and chimpanzee alleles are both present together in the shared transcriptional environment of allotetraploid piNs. In this study, unless otherwise indicated, when we describe gene expression differences (species bias) between humans and chimpanzees, we are referring to *cis*-regulated differences.

### Species-biased genes regulate excitatory neuron maturation and are mutated in developmental brain disorders

In the unstimulated condition, we found 1787 gene expression differences (FC>1.5, p-adj<0.05) between human and chimpanzee piNs (Figure 1d). These species-biased genes are enriched in pathways, including mitochondrial biology, that have previously been implicated in neuronal maturation (Figure 1e; Figure S2f) (Iwata et al., 2023). In addition, many of the species-biased genes that we identified control synaptic function (Koopmans et al., 2019), and pathogenic variants in some of these genes have been shown to cause neurodevelopmental disorders (NDDs). Fifty-three species-biased genes are known risk genes for autism spectrum disorder (ASD), including five genes classified as highest confidence ASD genes (Abrahams et al., 2013), underscoring that pathways that have evolved selectively in human cortical neurons are in some cases dysregulated in developmental brain disorders.

Notably, intersecting species-biased genes with genes that significantly change during early maturation in human layer II/III neurons *in vivo* shows significant enrichment for human-biased genes in the set of genes that decrease expression with maturation and significant enrichment for chimpanzee-biased genes in genes that increase expression with maturation (Figure 1f) (Herring et al., 2022). This suggests that many of the species-biased genes that we detected in tetraploid piNs are expressed during human and chimpanzee brain development, are likely associated with cortical neuron maturity, and may contribute to the prolonged process of human brain maturation. Our finding that maturation-associated genes are regulated in *cis* in cortical excitatory neurons underscores the cell-intrinsic, genetic underpinnings of slowed human neuronal maturation.

### A subset of activity-dependent genes are differentially expressed between human and chimpanzee piNs

Our initial goal was to determine if special features of activity-dependent human brain function are due to changes in activity-dependent genes, and whether these include changes to the expression of master regulator IEG TFs, the expression of downstream targets, or both. Towards this end, we depolarized piNs with KCl for 2hr to capture the expression of IEGs and 6hr to capture expression of LRGs. After 2 and 6 hr of depolarization, the activity-dependent changes in gene expression were highly correlated between diploid and tetraploid neurons and all the lines expressed canonical IEGs (Figure 2a; Figure S3a-d). From all samples, we identified 2778 ADGs at 2hr and 6580 ADGs at 6hr of exposure to KCl.

**Figure 2:**
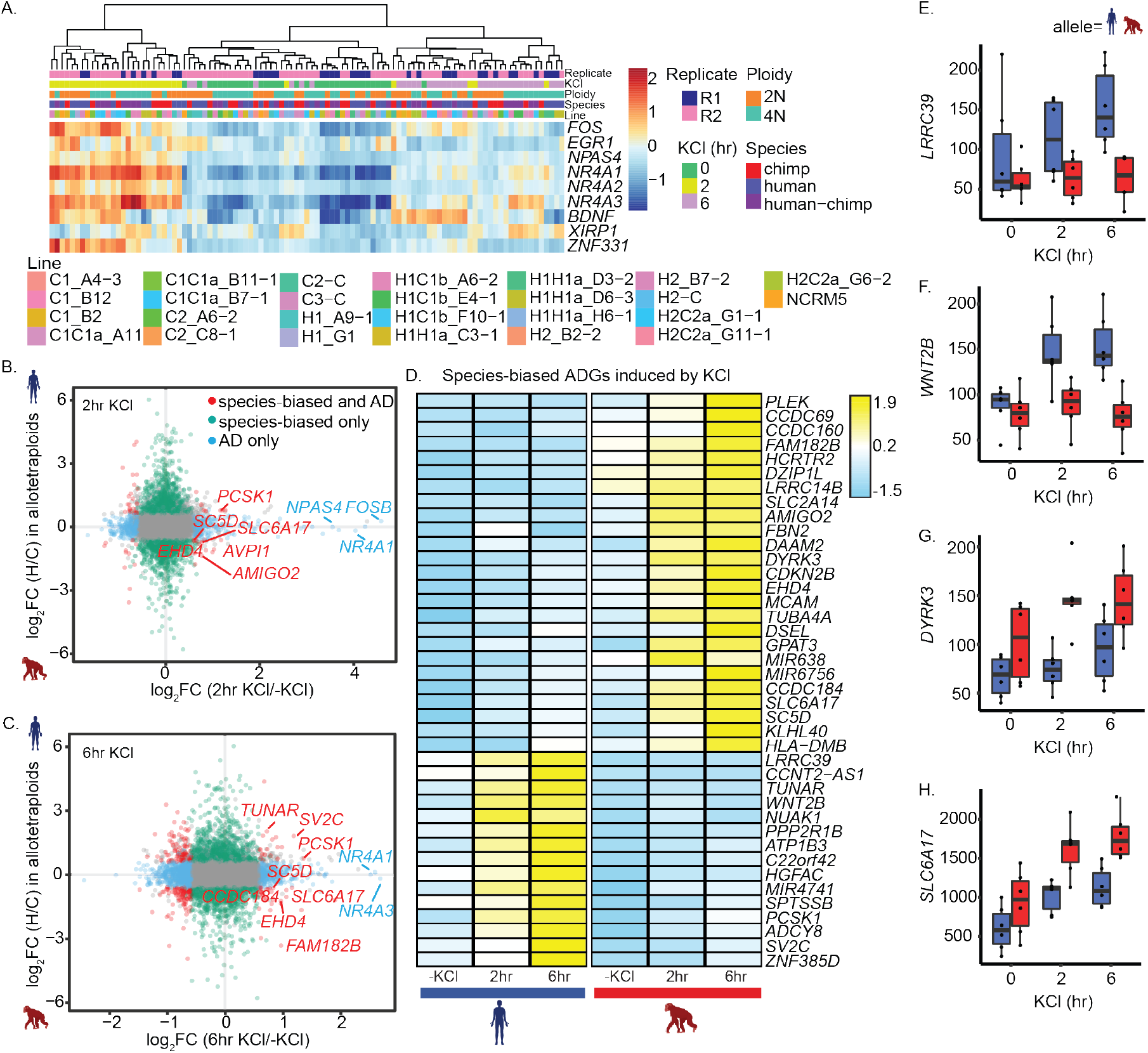
Identifying species-biased neuronal activity-dependent genes in piNs. a. Heatmap showing normalized expression values from RNA-seq for canonical ADGs in unstimulated and depolarized piNs. b. Scatterplot showing the log_2_ fold change in allelic expression (human/chimpanzee) vs. the log_2_ fold change in expression with 2hr KCl treatment. Red points indicate species-biased activity dependent genes, blue points indicate genes that are activity-dependent only, green points indicate genes that are species-biased only, and gray points are non-significant. c. Same as in b for 6hr KCl treatment. d. Normalized expression of species-biased activity-dependent genes that are upregulated with KCl treatment from allotetraploid piNs. Expression from human and chimpanzee alleles for all allotetraploids is averaged for each timepoint. e. Normalized expression at 0, 2, and 6hr KCl treatment for *LRRC39*, which is human-biased and activity-dependent. Each point shows the expression from the human or chimpanzee alleles in an individual allotetraploid piN line. Expression from human alleles is in blue and expression from chimpanzee alleles is in red. f. Same as in e for *WNT2B*. g. Same as in e for *DYRK3*, which is higher in chimpanzee. h. Same as in e for *SLC6A17*, which is higher in chimpanzee.

Although most activity-dependent genes were not differentially expressed between human and chimpanzee piNs, we identified 56 species-biased ADGs after 2hr of KCl exposure (Figure 2b) and 206 species-biased ADGs at 6hr after KCl treatment (Figure 2c). Analyzing our human samples alone, we did not identify any human-specific genes that are not present in the chimpanzee genome that were activity-dependent. None of the highly induced canonical IEG TFs, including *FOS, NPAS4*, and *NR4A1*, showed species-specific differences in expression between human and chimpanzee piNs, suggesting there is not a global up- or down-regulation of activity-dependent programs across species.

We identified a subset of species-biased activity-dependent genes that are activity-dependent only in one species, suggesting that they may have acquired new regulatory regions that specifically control the response to stimulus through evolution (Figure 2d-h). These include the human-biased genes *WNT2B, LRRC39*, and *TUNAR* that are highly induced in human piNs, but not in chimpanzee piNs across the stimulation time course (Figure 2d-f). We also identified species-biased activity-dependent genes, including *SV2C, SLC6A17*, and *SC5D*, that are induced by activity to a similar degree in both species, but have overall lower expression levels in one species compared to the other (Figure 2d,g,h). Taken together, this suggests that there are evolutionary changes to the ADGE program that occur both by making some genes more stimulus-responsive while changing the overall expression of other genes.

A subset of the species-biased activity-dependent genes are known to control synaptic function and be mutated in neurodevelopmental disorders. After 2hr and 6hr of KCl treatment, there are ten species-biased activity-dependent genes that are known to act at the synapse (*ADCY8, CHRNA3, CHRNB4, KCTD12, NRP1, PPP2R1B, SLC18A3, SLC6A17, SV2C*) (Koopmans et al., 2019). *SHISA9* is upregulated in human and chimpanzee piNs in response to 2hr of depolarization, but its expression levels are higher at all timepoints in human piNs. SHISA9 modulates synaptic AMPA receptors, and the *SHISA9* locus has been implicated in schizophrenia (Liu et al., 2021). *CEP19, DPH5, H4C9, SC5D, BBS12, ERF*, and *SLC6A17* are all classified as high-confidence neurodevelopmental disorder (NDD) genes (Abrahams et al., 2013; Leblond et al., 2021). *SLC6A17* is an activity-dependent gene that is more highly expressed in chimpanzee piNs than in human piNs. *SLC6A17* encodes a solute carrier involved in the presynaptic uptake of neurotransmitters, and variants within *SLC6A17* are associated with intellectual disability (Iqbal et al., 2015). Notably, a prior study demonstrated that *SLC6A17* is a marker of piN maturation (Shan et al., 2024). The increased expression of *SLC6A17* in chimpanzee piNs raises the possibility that SLC6A17 contributes to the more rapid maturation of chimpanzee neurons via its role at the synapse. *NUAK1* encodes an activity-dependent kinase that is induced to a higher level in human compared to chimpanzee piNs. NUAK1 promotes axon growth and branching, as well as synaptic maturation by increasing mitochondrial activity in the axon (Courchet et al., 2013; Lanfranchi et al., 2024). Thus, it may be playing a role in increasing human neuron morphological complexity and regulating neuronal maturation (Iwata et al., 2023). Taken together, these findings suggest that the ADGE program has changed between human and chimpanzee neurons by fine-tuning the expression of many LRGs that may then impact the myriad cellular functions associated with neuronal maturation.

### Identifying species-specific open chromatin regions in response to neuronal activity

To understand how these human- and chimpanzee-biased features of ADGE evolved, we turned our attention towards defining the gene regulatory elements (e.g enhancers and promoters) that control the expression of species-biased ADGs, as sequence variants in gene regulatory elements are thought to be major contributors in evolution (King and Wilson, 1975; Carroll, 2008; McLean et al., 2011). Towards this end, we used ATAC-seq, which identifies accessible genomic regions, as a first step towards identifying enhancers and promoters that are differentially activated between membrane-depolarized human and chimpanzee piNs (Buenrostro et al., 2013). We performed ATAC-seq using a subset of the diploid, autotetraploid, and allotetraploid piNs that were used for our RNA-seq experiments. We left the piNs untreated or exposed them to an elevated level of KCl for 30 minutes or 2hr to capture changes in chromatin accessibility that precede changes in IEG and LRG expression. Principal component analysis showed that our ATAC-seq samples separated along the first two PCs by species as expected, with human-chimpanzee tetraploid samples falling between human and chimpanzee diploid and autotetraploids (Figure 3a). In addition, we used CUT&Tag (Kaya-Okur et al., 2019) in allotetraploid piNs to identify sites across the genome with the histone modification H3K27Ac, a mark that is found at active enhancers (Creyghton et al., 2010).

**Figure 3:**
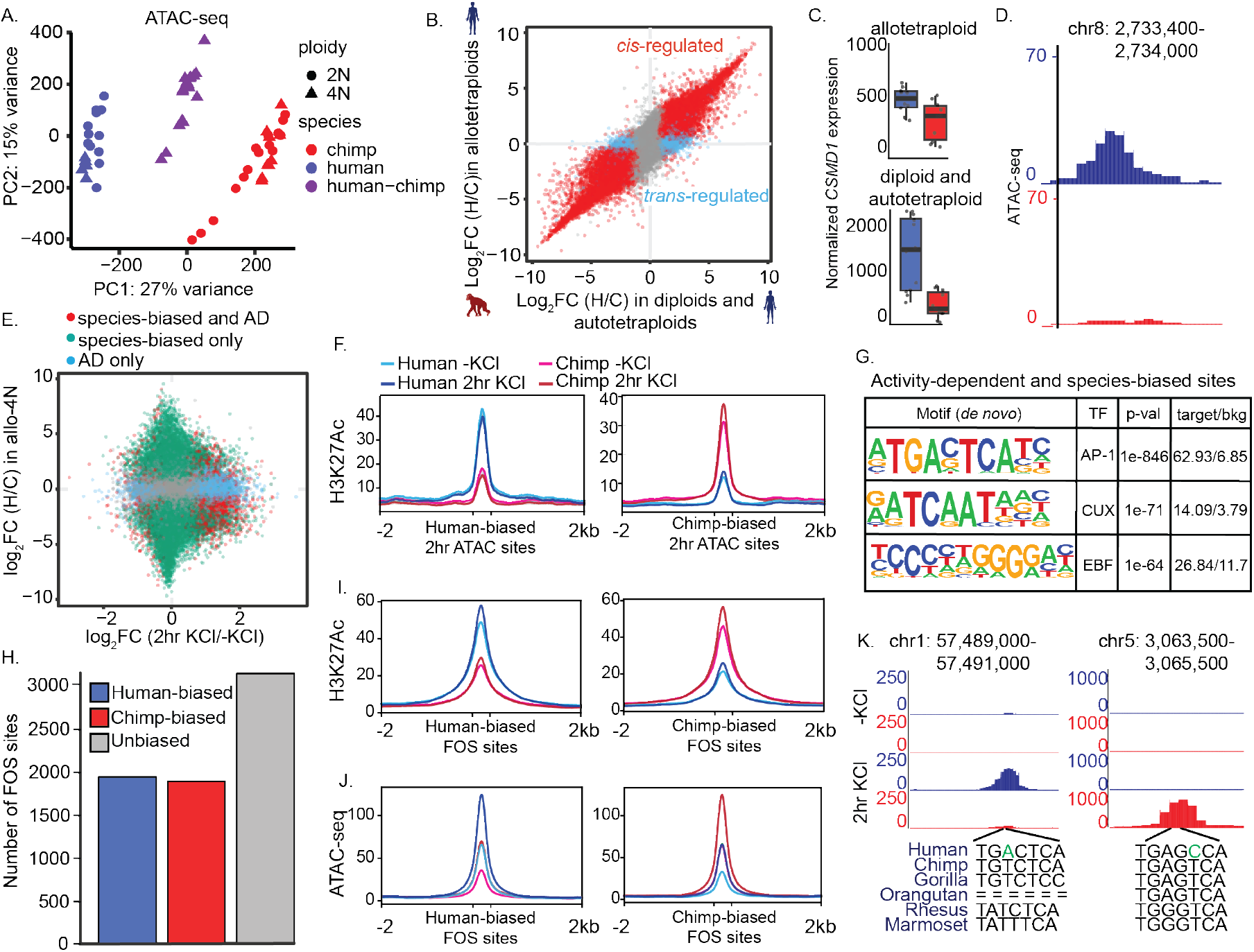
Species-biased activity-dependent open chromatin regions in piNs are enriched for FOS binding sites. a. PCA of ATAC-seq data from diploid and tetraploid piNs showing PC1 vs. PC2. b. Scatterplot of ATAC-seq data showing the log_2_ fold change in accessibility between chimpanzee and human alleles in allotetraploid cells vs. the log_2_ fold change in accessibility between chimpanzee and human diploid and autotetraploid piNs. *Cis*-regulated sites are indicated in red and *trans*-regulated sites are indicated in blue. c. Expression of the species-biased gene *CSMD1* in allotetraploid (top) and diploid and autotetraploid (bottom) piNs. Blue bars show expression from human alleles and red bars show expression from chimpanzee. d. Tracks showing a species-biased accessible site near *CSMD1*. e. Scatterplot showing the log_2_ fold change in allelic accessibility (human/chimpanzee) vs. the log_2_ fold change in accessibility with KCl treatment (2hr KCl/-KCl) from ATAC-seq data. Red points indicate species-biased and activity-dependent (AD) sites, blue points indicate sites that are activity-dependent only, green points indicate sites that are species-biased only, and gray points are non-significant. f. Average diagram of H3K27Ac CUT&Tag signal at human-biased (left) and chimpanzee-biased (right) activity-dependent ATAC-seq regions. CUT&Tag data is from three allotetraploid piN lines and the signals from human and chimpanzee alleles are separated in the plot. g. *De novo* motif enrichment analysis at activity-dependent and species-biased ATAC-seq peaks after 2hr KCl treatment. h. Bar plot showing the number of activity-dependent FOS binding sites from allotetraploids that are human-biased, chimpanzee-biased, and unchanged between species. i. Average diagram of H3K27Ac CUT&Tag signal at human-biased and chimpanzee-biased activity-dependent FOS binding sites. CUT&Tag data is from allotetraploids and signals from human and chimpanzee alleles are separated in the plot. j. Same as in e for ATAC-seq signal. k. Example of a human-biased (left) and chimpanzee-biased (right) FOS binding site. Tracks are separated for human and chimpanzee alleles with and without KCl from allotetraploid piN lines. Sequences of the AP-1 motifs within these FOS binding sites are indicated for human, chimpanzee, gorilla, orangutan, rhesus, and marmoset from UCSC and derived SNVs in the human are indicated in green.

Within the piN ATAC-seq samples, we identified 44,789 regions of open chromatin. Among the open chromatin regions that we detected in the uninduced tetraploid piNs, 11,191 displayed a significant difference in accessibility between human and chimpanzee piNs (Figure 3b). Of these, 1669 are located close to a gene (532 genes in total) that displayed a human or chimpanzee bias in expression. This raises the possibility that these regions correspond to enhancers or promoters that differentially regulate gene expression between species. For example, *CSMD1*, an ASD-associated gene (Abrahams et al., 2013) that regulates the complement pathway in mouse and human neurons and plays a role in synapse elimination during development (Baum et al., 2024), has human-biased expression and is near multiple human-biased open chromatin regions (Figure 3c,d).

To begin to identify activity-dependent regulatory regions that are species-biased or differentially active in human and chimpanzee piNs, we compared the open chromatin and H3K27Ac signal before and after membrane depolarization. At 30 min after stimulation, we identified 575 activity-dependent accessible regions. Of these, 102 are species-biased between human and chimpanzee piNs (Figure S4a). Following 2hr of membrane depolarization, we identified 8699 activity-dependent accessible regions and of these, 1845 displayed a bias in their accessibility between human and chimpanzee piNs (Figure 3e). Human-biased accessible regions displayed human-biased H3K27Ac signal and chimpanzee-biased accessible regions displayed chimpanzee-biased H3K27Ac signal (Figure 3f). Altogether, among the species biased activity-dependent open chromatin regions, we identified 37 that were close to 29 species-biased activity-dependent genes, raising the possibility that these regions may contribute to the species-biased expression of these nearby genes.

### Species-biased activity-dependent open chromatin sites are highly enriched for AP-1 motifs

The differential accessibility of activity-dependent open chromatin regions in human and chimpanzee piNs may be due to DNA sequence differences between humans and chimpanzees that then give rise to species-specific differences in TF binding. To identify potential TF binding sites within open chromatin regions that display a bias in human and chimpanzee piNs, we performed motif enrichment analysis for species-biased open chromatin regions. We found that the species-biased open chromatin regions present at 30 min after membrane depolarization were enriched for CUX TF family motifs (22% of sites, p=1e-15), which are known to regulate genes that specify the differentiation of cortical neurons (Cubelos et al., 2010), and CREB binding motifs (19.6% of sites, p=1e-11), which are known to regulate activity-dependent promoter activation in neurons (Sheng et al., 1988; Sheng and Greenberg, 1990) (Figure S4b). By contrast, the species-biased sites that we identified after 2hr of membrane depolarization were highly enriched for motifs for the AP-1 TFs, including FOS and JUN family members (63% of species-biased AD sites, vs 7% of genomic background, p=1e-846) (Figure 3g). This enrichment was much stronger than the next highest enrichment of CUX TF motifs at only 20% of species-biased AD sites. FOS is a classic marker for activated cells in the brain and has more recently been shown to be critical for learning and memory in mice (Paylor et al., 1994; He et al., 2002; Countryman et al., 2005; Yap et al., 2021). FOS binds enhancers as a heterodimer with JUN TFs, which recruits the chromatin remodeling complex BAF in response to stimuli, leading to the eviction of histones and an increase in chromatin accessibility (Vierbuchen et al., 2017; Stroud et al., 2020). Given the remarkable enrichment of AP-1 motifs within regions of open chromatin that are differentially active in human and chimpanzee piNs, we next asked whether FOS binds to these regions in a species-specific manner in response to neuronal activity.

### FOS binding is evolutionarily dynamic in excitatory neurons

We profiled FOS binding in three allotetraploid piN lines before and after membrane depolarization using CUT&Tag. The vast majority of FOS binding was detected following 2hr of exposure to KCl, which is consistent with FOS being largely absent in uninduced piNs (Table S4). We identified 6973 inducible FOS binding sites. Remarkably, 1940 of the sites bound significantly more FOS on the human allele than the chimpanzee allele (human-biased sites), and 1892 bound significantly more FOS on the chimpanzee allele than the human allele (chimpanzee-biased sites) (Figure 3h). We were surprised to see that over half of the FOS binding sites displayed differential binding between the human and chimpanzee alleles, with a similar proportion human-biased as chimpanzee-biased, suggesting that FOS binding is highly dynamic during evolution. The human-biased FOS binding sites displayed a similar bias in inducible H3K27Ac and ATAC-seq signal after 2hr of depolarization, and vice versa for chimpanzee-biased FOS sites (Figure 3i,j). Notably, unbiased FOS binding sites have similar levels of H3K27Ac and accessibility on human and chimpanzee alleles (Figure S4c,d).

We hypothesized that species-biased FOS binding sites may regulate species-biased activity-dependent genes. For eight species-biased FOS binding sites, the nearest gene was species-biased and activity-dependent after 2hr KCl treatment, and for 32 species-biased FOS binding sites, the nearest gene was species-biased and activity-dependent after 6hr KCl treatment (near 11.2% of species-biased ADGs vs. 8.4% near all expressed genes, p=0.07). These include binding sites near the cis-regulated genes *TUNAR* and *PCSK1*, which we discuss further below.

### Species-biased FOS binding sites are enriched for SNVs in AP-1 motifs

To understand the evolution of species-biased FOS binding sites, we next asked if SNVs within AP-1 sites between humans and chimpanzees cause species-biased FOS binding (Figure 3k). FOS-bound AP-1 sites in neurons are generally highly conserved between species and constrained within the human population (Greenberg Lab, unpublished observation), suggesting that evolutionary changes to these sequences may have functional consequences. Remarkably, we found that within human-biased and chimpanzee-biased FOS binding sites, AP-1 motifs are highly enriched for SNVs compared to AP-1 sites in binding sites that do not display a species bias (Figure 4a). Within the human-biased sites, there are 323 variants within the 2135 AP-1 motifs (variant frequency = 0.021). Of these, 153 variants are newly derived in the human lineage and create a complete AP-1 motif that is not present in the chimpanzee, gorilla, macaque, or marmoset genomes (example in Figure 3k). In chimpanzee-biased sites, there are 311 variants within 2085 AP-1 motifs (variant frequency = 0.021). Of these, 259 variants are newly derived in the human lineage and result in a disruption of the AP-1 sequence motif in humans but not in other primates. By contrast, there are only 39 SNVs within the 3271 AP-1 motifs in unbiased sites (variant frequency = 0.0017, chi-squared p-val human-biased vs. unchanged and chimpanzee-biased vs. unchanged <2.2e-16) (Figure 4a).

**Figure 4:**
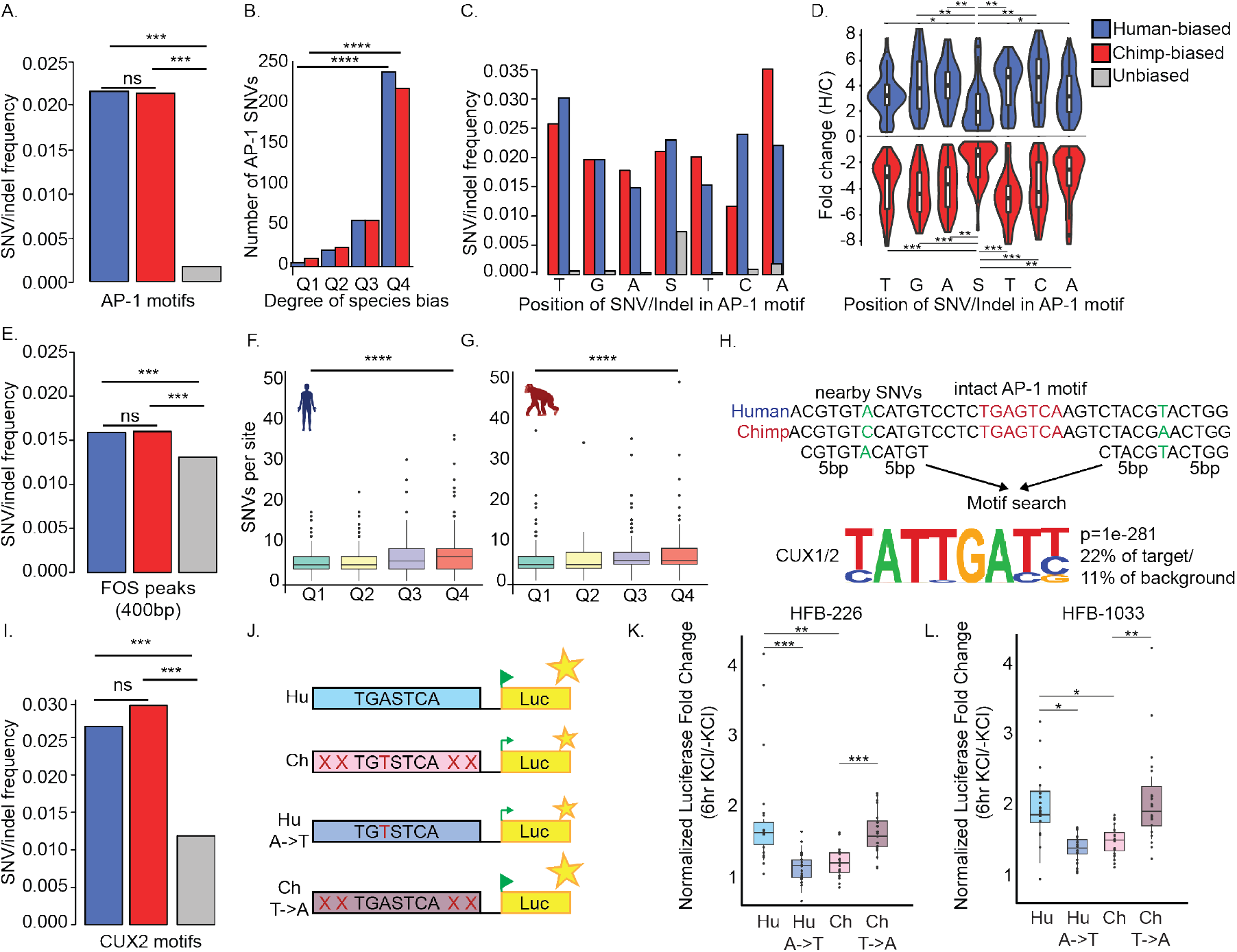
SNVs within FOS binding sites drive species differences in enhancer activity. a. The SNV and indel frequency within AP-1 motifs in human-biased, chimpanzee-biased, and unbiased FOS binding sites. For all panels, blue bars are human-biased sites, red bars are chimpanzee-biased sites, and gray bars are unbiased sites. Chi-squared p-val human-biased vs. unbiased and chimpanzee-biased vs. unbiased p <2.2e-16. b. Barplot showing the number of SNVs and indels within AP-1 motifs in quartiles 1-4 of human-biased and chimpanzee-biased FOS binding sites. Q1 has the lowest absolute fold change difference in binding between species, and Q4 has the highest absolute fold change difference (See Supplemental Figure 5a,b). c. Barplot showing the SNV/indel frequency at each position within the AP-1 motif at human-biased, chimpanzee-biased, and unbiased FOS binding sites. “S” in the central position is either a “G” or “C”. d. Violin plot showing the bias in FOS binding (Human allele/Chimpanzee allele [H/C]) for human-biased and chimpanzee-biased sites that have SNVs or indels at each of the AP-1 motif positions. KS test for significance with BH correction comparing position 4 in the AP-1 motif to each other position. e. Barplot showing the SNV/indel frequency within the entire 400bp FOS binding region for human-biased, chimpanzee-biased, and unbiased sites. f. Boxplot showing the number of SNVs per FOS binding site for human-biased sites in quartiles 1-4. g. Same as in j for chimpanzee-biased sites. h. Diagram of the motif search methodology for SNVs in human-biased and chimpanzee-biased FOS binding sites outside of AP-1 motifs. i. Barplot showing the SNV/indel frequency within CUX2 motifs in human-biased, chimpanzee-biased, and unbiased FOS sites. j. Diagram showing the AP-1 SNV sequence change experiments for luciferase assays. HFB-226 and HFB-1033 have multiple SNVs throughout the 400bp binding region, but only the AP-1 SNVs in positions 3 or 5 were changed in this assay. “X”s in the construct signify other SNVs between human and chimpanzee within the FOS-bound region. k. Boxplot showing luciferase results for HFB-226 when the AP-1 SNV is changed between chimpanzee and human (* p < 0.05, ** p < 0.01, *** p < 0.001; Bonferroni corrected p-values from Wilcoxon Rank Sum Test). l. Same as in k for HFB-1033.

We divided species-biased FOS binding sites into quartiles, with Q4 having the greatest species bias in FOS binding and Q1 having the smallest species bias, and asked whether the presence of AP-1 SNVs was associated with a larger species bias in binding (Figure S5a,b). Human-biased, chimpanzee-biased, and unbiased sites in all four quartiles were found to have a similar level of inducible FOS binding upon KCl stimulation (Figure S5c). We found that Q4 sites have the most AP-1 SNVs (49.5% of human-biased Q4 sites have AP-1 SNVs vs 1% of human-biased Q1 sites, chi-squared p-val<0.00001; 47% of chimpanzee-biased Q4 sites have AP-1 SNVs vs. 2% of Q1 sites, chi-squared p-val<0.00001) (Figure 4b), suggesting that disruption of the core AP-1 motif has a strong effect on the strength of FOS binding.

Prior work found that within the seven nucleotide AP-1 motif sequence, changes to the A and T in the third and fifth positions had the greatest effect on FOS binding. Changes in positions 1, 2, 6, and 7 had an intermediate effect, and the central base had relatively little effect on FOS binding (Yang et al., 2022). When we examined which bases within the AP-1 motif change most frequently in species-biased sites, we found examples of changes at each of the seven AP-1 bases (Figure 4c). In contrast, the central base, which is not critical for FOS binding, is most frequently changed in AP-1 sites that display no bias in FOS binding in human and chimpanzees piNs. Notably, SNVs are rarely present at the other six nucleotides within the AP-1 motif of unbiased FOS binding sites (Figure 4c). Consistent with the weak effect of the central base on FOS binding, the fold change in FOS binding is lower at species-biased sites with an SNV at the central base compared to the fold change when other bases within the AP-1 site contain an SNV (Figure 4d). Taken together, these findings strongly suggest that evolutionary changes at core bases within AP-1 sites drive human-specific activity-dependent FOS binding to enhancers.

### Sequence changes outside of AP-1 motifs affect FOS binding

Though AP-1 motifs within species-biased FOS binding sites are enriched for SNVs compared to unbiased sites, many of the AP-1 motifs in human-biased and chimpanzee-biased FOS binding sites have no SNV. This suggests that SNVs outside of AP-1 motifs, but within the broader FOS-bound genomic region, may also affect FOS binding. Prior work showed that SNVs in bases very close to AP-1 motifs, or in the motifs of cooperating lineage-specific TFs, can affect FOS binding (Yang et al., 2022). We found that SNV frequency within the 400 bp peak region for each FOS binding site is significantly higher in species-biased compared to unbiased sites (variant frequency =0.016 in species-biased sites, 0.013 in unbiased sites, chi-squared p-val human-biased vs. unbiased and chimpanzee-biased vs. unbiased <2.2e-16) (Figure 4e). Notably, this frequency is still significantly lower than the SNV frequency within AP-1 motifs at species-biased sites, reinforcing the importance of AP-1 sequence fidelity for FOS binding (AP-1 motifs vs. peak region in human-biased sites chi-squared p-val<2.2e-16; in chimpanzee-biased sites chi-squared p-val= 2.1e-07) (Figure 4e,a). Sites in Q4, which have the greatest bias in binding between species, have an increased number of SNVs outside of AP-1 sites compared to sites in quartile 1 (human-biased sites p-val=5.11e-16, chimpanzee-biased sites p-val=2.1e-9, Wilcoxon rank sum test) (Figure 4f,g), suggesting that a greater number of SNVs between species gives rise to larger magnitude changes in FOS binding.

We sought to identify TFs other than FOS where an SNV in their binding site might secondarily affect FOS binding at species-biased FOS sites. We searched for TF motifs in a 10bp window around all SNVs detected within human- or chimpanzee-biased FOS binding sites that did not fall within an AP-1 motif (Figure 4h). Motif enrichment analysis at these non-AP-1 motif SNVs revealed that these regions are enriched for CUX and EBF TF family motifs (Figure 4h). For peaks in which CUX and AP-1 motifs are both found, they were a median of 60bp apart (Figure S5d). CUX family TFs, CUX1 and CUX2, have been used as markers for cortical excitatory neurons in layer II/III in the mouse (Nieto et al., 2004). Both CUX1 and CUX2 bind to and regulate genes that control dendritic outgrowth and dendritic spine and synapse formation (Cubelos et al., 2010). We found that within species-biased FOS binding sites, CUX motifs are enriched for SNVs compared to the surrounding peak region (Figure 4i,e) (CUX2 motifs vs. peak region in human-biased and chimpanzee-biased sites chi-squared p-val<2.2e-16). In addition, CUX motifs in species-biased FOS sites have a higher SNV frequency than FOS-bound sites that are unbiased (human-biased vs unbiased sites, chi-squared p-val = 1.6e-11; chimpanzee-biased vs. unbiased sites, chi-squared p-val = 2.3e-14) (Figure 4i). CUX motifs were recently identified in another FOS binding dataset from piNs, where it was suggested that CUX factors may cooperate with FOS to activate enhancers during differentiation (Sanchez-Priego et al., 2022). Our data suggest that FOS and CUX1/2 may not only bind together at enhancers, but that SNVs in either one of their motifs may modulate enhancer activity during human evolution.

### Changes in AP-1 motif SNVs drive species-specific AD enhancer activity in neurons

To determine if human-specific changes within FOS-bound AP-1 sites affect enhancer activity, we tested the effects of human-specific variants using a luciferase reporter assay. Given that luciferase assays often cannot recapitulate *in vivo* enhancer activity, we first screened 19 species-biased FOS binding sites to identify elements that are able to drive activity-dependent luciferase expression in reporter assays (Figure S5e,f). We chose human-biased FOS sites that have an SNV in the 3rd or 5th position of the AP-1 motif that is derived in the human lineage. We cloned these species-biased FOS binding sites (500 bp-1 kb) upstream of a nanoluciferase gene with a minimal promoter, transfected the constructs into cortical neurons dissociated from embryonic mouse brains, and performed luciferase assays before and six hours after membrane depolarization. We identified two human-biased FOS binding sites (HBF), HBF-226 and HBF-1033, that were highly effective at driving luciferase expression upon membrane depolarization (Figure S5e). When compared with the chimpanzee version of the same binding site, we found that the inducibility was significantly greater for the human sequence for both HBF-226 and HBF-1033, consistent with our finding that both of these regions display increased FOS binding to the human compared to the chimpanzee allele in tetraploid piNs (Figure S5g,h).

To determine whether the SNV within the AP-1 motif at these two sites causes differential enhancer activity between human and chimpanzee piNs, we swapped the human and chimpanzee SNVs in the AP-1 motif in HBF-226 and HBF-1033, while leaving all other SNVs in the 500bp region unchanged (Figure 4j). For HBF-226, we changed the A in position 3 of the AP-1 motif to the chimpanzee T within the human construct and vice versa (Figure S5i). For HBF-1033, we changed the T in position 5 of the AP-1 motif to the chimpanzee C in the human construct and vice versa (Figure S5j). For both HBF-226 and HBF-1033, swapping the human AP-1 SNV into the chimpanzee sequence resulted in a significant increase in the inducible luciferase activity to the same level as that observed from the human sequence (Figure 4k,l). This suggests that lone SNVs within AP-1 motifs in humans are sufficient to convert the relatively inactive chimpanzee sequences into effective drivers of activity-dependent reporter gene expression. Similarly, putting the chimpanzee SNV into the human sequence significantly reduced enhancer activity in both HBF-226 and HBF-1033 (Figure 4k,l), suggesting that these SNVs in the AP-1 motif are also necessary for enhancer activity in this context. The effect of this single base change is striking, as there are 11 other SNVs in HBF-226 and 13 other SNVs and 4 indels in HBF-1033. These results suggest that single base changes in AP-1 motifs during evolution can have strong effects on enhancer function.

### Species-biased FOS binding sites are bona fide enhancers of ADGs

Luciferase assays can measure the enhancer activity of a DNA sequence outside of the context of chromatin, but do not reveal whether a given region is an enhancer for a specific gene. Testing the activity of an enhancer within its genomic context and identifying its *in vivo* target genes has historically been challenging, but CRISPR-inhibition (CRISPRi) technology now makes it possible to direct chromatin silencing to enhancer loci and measure the effect on gene expression. To test whether the evolution of a new activity-dependent FOS binding site in human neurons led to increased expression of a species-biased activity-dependent gene, we repressed FOS binding sites within their genomic context. We used CRISPRi to target the human-biased FOS binding sites near the human-biased activity-dependent genes *TUNAR* and *PCSK1. TUNAR* encodes a conserved microprotein pTUNAR, which in the mouse plays a role in neuronal development by regulating intracellular calcium dynamics (Senís et al., 2021). *PCSK1* encodes proprotein convertase 1, which cleaves neuropeptides in the brain, including the FOS target SCG2 (Fischer-Colbrie et al., 1995).

We identified two FOS-bound regions 13 kb downstream of *TUNAR* (Figure 5a), a gene that is upregulated 3-fold from human alleles with depolarization, but minimally from chimpanzee alleles (Figure 5b). We named these regions putative enhancer 1 (*TUNAR*-PE1) and 2 (*TUNAR*-PE2). FOS binds to these regions in an activity-dependent manner, with stronger binding to human alleles compared to chimpanzee alleles. These regions also have human-biased activity-dependent H3K27Ac and ATAC-seq signals, suggesting that they may act as human-biased activity-dependent enhancers (Figure 5a).

**Figure 5:**
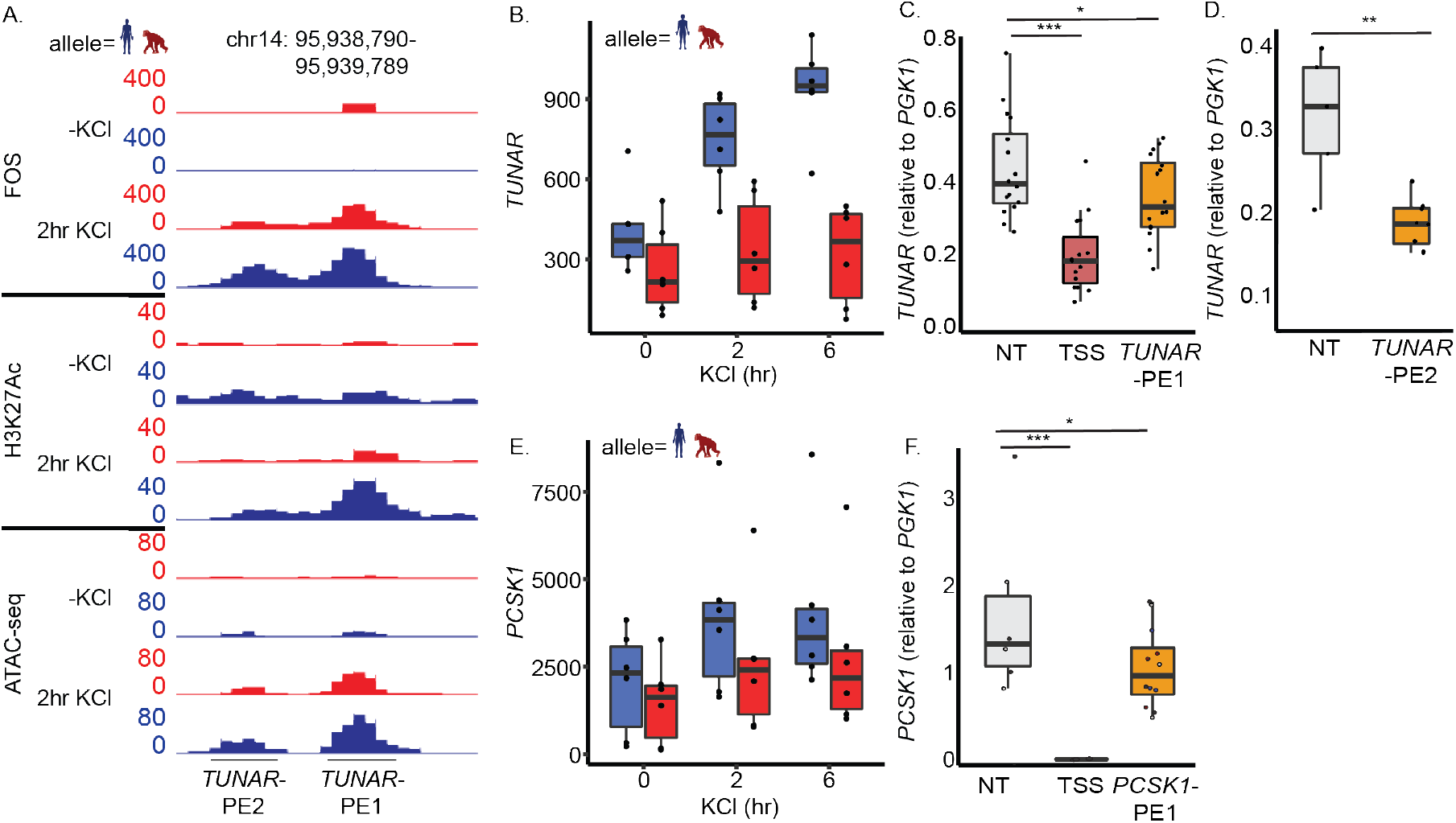
FOS binding sites near *TUNAR* and *PCSK1* are activity-dependent enhancers for these genes. a. Tracks from the UCSC genome browser showing the putative enhancers, PE1 and PE2, downstream of the *TUNAR* gene. Tracks are from one human-chimpanzee allotetraploid cell line H2C2a G6-2, and show FOS CUT&Tag, H3K27Ac CUT&Tag, and ATAC-seq at -KCl and 2hr KCl timepoints. Reads from human alleles are in blue and reads from chimpanzee alleles are in red. b. Boxplot showing the expression of *TUNAR* from human-chimpanzee allotetraploid piNs at 3 timepoints: 0, 2, and 6 hours KCl. Reads from the human and chimpanzee alleles are separated. Expression from human alleles is in blue and expression from chimpanzee alleles is in red. c. Boxplots of qPCR data showing expression of *TUNAR* after 6hr KCl treatment from CRISPRi cells when using nontargeting sgRNAs (NT), sgRNAs targeting *TUNAR* TSS (TSS), and sgRNAs targeting *TUNAR*-PE1 (*TUNAR*-PE1). d. Same as in c for *TUNAR*-PE2. e. Boxplot showing the expression of *PCSK1* from human-chimpanzee allotetraploid piNs at 3 timepoints: 0, 2, and 6 hours KCl. Reads from the human and chimpanzee alleles are separated. Expression from human alleles is in blue and expression from chimpanzee alleles is in red. f. Boxplots of qPCR data showing expression of *PCSK1* from CRISPRi cells when using nontargeting sgRNAs (NT), sgRNAs targeting *PCSK1* TSS (TSS), and sgRNAs targeting *PCSK1*-PE1 (*PCSK1*-PE1).

To test whether these elements function as bona fide activity-dependent enhancers for *TUNAR*, we generated stable hiPSC lines that constitutively express dCas9-KRAB as well as a synthetic guide RNA (sgRNA) targeting one of the two PEs. We targeted the transcription start site (TSS) as a positive control. As a negative control, we generated lines expressing two different non-targeting (NT) sgRNAs (Tian et al., 2019). As expected, targeting the TSS of *TUNAR* with each of two different sgRNAs significantly knocked down *TUNAR* expression (Figure 5c). At *TUNAR*-PE1 and *TUNAR*-PE2, we targeted sgRNAs close to the AP-1 site, based on the assumption that blocking FOS binding to DNA would have the largest effect on potential enhancer activity (Figure S6a,b). When we targeted *TUNAR*-PE1 with either of two sgRNAs, we observed a significant reduction in activity-dependent *TUNAR* expression by qPCR when compared to NT controls. (Figure 5c). When we targeted *TUNAR*-PE2 with either of three sgRNAs, we saw an even greater and significant reduction in activity-dependent *TUNAR* expression (Figure 5d). Results were consistent between two hiPSC lines, NCRM5 and H2-C. Together these findings suggest that both *TUNAR*-PE1 and *TUNAR*-PE2 function as enhancers for the gene *TUNAR*, with *TUNAR*-PE2 being a stronger enhancer.

We also identified a FOS binding site 130 kb downstream of *PCSK1*, which we term *PCSK1*-PE1, that is more highly bound on human alleles than chimpanzee alleles in piNs. *PCSK1* is upregulated in response to depolarization in human and chimpanzee piNs, but is more highly expressed at all time points from human alleles (Figure 5e). As expected, targeting the TSS of *PCSK1* with CRISPRi fully inhibited *PCSK1* expression (Figure 5e). When we targeted *PCSK1*-PE1 with each of four sgRNAs, activity-dependent *PCSK1* expression was significantly reduced (Figure 5f; Figure S6c). Together, these experiments demonstrate that these three human-biased FOS binding sites are bona fide enhancers for the human ADGs *TUNAR* and *PCSK1*.

### Human-biased activity-dependent genes affect neuronal firing dynamics

To begin to assess how human-specific changes in the evolution of ADGE affect neuronal firing, we investigated the function of three human-biased activity-dependent genes, *TUNAR, SV2C*, and *PCSK1*, using multielectrode arrays (MEAs) (Figure 6a). When co-cultured with human iPSC-derived astrocytes over 39 days, wild-type human piNs become increasingly electrically active and display an increase in synchronous firing across the neuronal culture (Figure 6b, Figure S7a,b), an indication that the neurons are maturing over time (Wagenaar et al., 2006; Napoli and Obeid, 2015; Fair et al., 2020).

**Figure 6:**
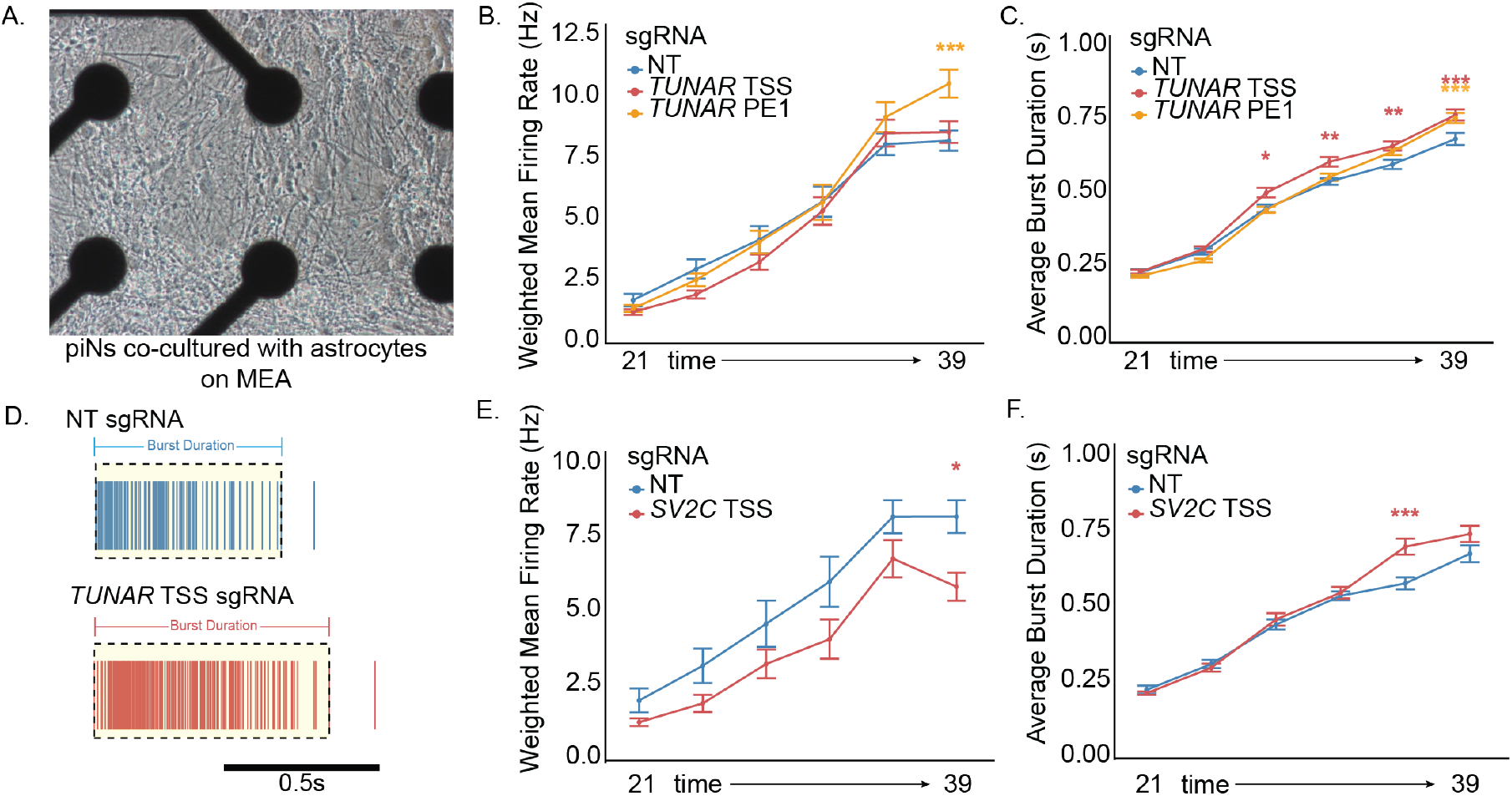
*TUNAR* or *SV2C* knockdown modulates neuronal firing in culture. a. Brightfield image of piNs and human iPSC-derived astrocytes cultured on 48-well multielectrode array (MEA) plate. b. Plot of weighted mean firing rate in *TUNAR* TSS, *TUNAR*-PE1, and NT CRISPRi lines recorded every 3/4 days from Day 21 to Day 39. Error bars are ± SEM (*** p < 0.001; Holm-Sidak corrected p-values from t-tests on estimated marginal means derived from mixed effects models). c. Plot of average burst duration in *TUNAR* TSS, *TUNAR*-PE1, and NT CRISPRi lines recorded every 3/4 days from Day 21 to Day 39. Error bars are ± SEM (* p < 0.05, ** p < 0.01, *** p < 0.001; Holm-Sidak corrected p-values from t-tests on estimated marginal means derived from mixed effects models). d. Raster plot of spikes recorded from NT CRISPRi and *TUNAR* TSS lines at Day 39 showing differences in burst duration. e. Plot of weighted mean firing rate in *SV2C* TSS and NT CRISPRi lines recorded every 3-4 days from Day 21 to Day 39. Error bars are ± SEM (* p < 0.05; Holm-Sidak corrected p-values from t-tests on estimated marginal means derived from mixed effects models). f. Plot of average burst duration in *SV2C* TSS and NT CRISPRi lines recorded every 3/4 days from Day 21 to Day 39. Error bars are ± SEM (*** p < 0.001; Holm-Sidak corrected p-values from t-tests on estimated marginal means derived from mixed effects models).

To investigate the function of *TUNAR* in human neurons, we cultured cell lines that express dCas9-KRAB with either sgRNAs targeting the *TUNAR* TSS or non-targeting control sgRNAs on MEAs, and compared the firing properties of the piNs. We found that at day 28 in culture, the neurons in which *TUNAR* expression is inhibited have differentiated normally and show no significant difference in gene expression when compared to cells expressing the non-targeting sgRNAs (Figure S7c). However, inhibition of *TUNAR* expression (Figure 5c) led to an increase in the duration of neuronal bursts during neuronal firing (Figure 6c,d). This change in bursting appeared early during the course of neuronal maturation, and increased as the neurons matured (Figure 6c). By contrast, inhibition of *TUNAR* expression did not affect the overall firing rate of the neurons (Figure 6b), although we did observe a small decrease in the number of bursts at some time points (Figure S7d). Overall, these effects are consistent with pTUNAR’s known role in calcium sequestration in the endoplasmic reticulum (Senís et al., 2021).

We also investigated the function of *SV2C*, a human-biased activity-dependent gene that encodes a synaptic vesicle protein (Figure 2c). Targeting the promoter of *SV2C* with a pool of sgRNAs led to a substantial decrease in *SV2C* expression compared to the non-targeting sgRNAs at day 28 in culture, but did not affect the expression of any other genes. This suggests that neurons differentiate normally when *SV2C* expression is significantly reduced (Figure S7e). However, *SV2C* appears to control neuronal firing. We found that *SV2C* knockdown leads to an overall decrease in the weighted mean firing rate of the neurons (Figure 6e), and that as the neurons mature, *SV2C* knockdown leads to an increase in burst duration and a higher frequency of spiking during bursting (Figure 6f, Figure S7f). Within the SV2C amino acid sequence, there are two binding sites for synaptotagmin: one that facilitates vesicle release and another that inhibits vesicle release under high Ca2+ conditions (Schivell et al., 2005; Dardou et al., 2013; Dunn et al., 2017), suggesting possible mechanisms by which the decrease in *SV2C* expression affects neuron firing and bursting dynamics. In contrast to our finding that a decrease in the expression of the human-biased genes *TUNAR* and *SV2C* affects neuronal firing properties, we found that inhibiting the expression of *PCSK1* had no significant effect on neuronal firing (Figure S7g,h).

Finally, to determine if human-specific changes in a single enhancer region can affect neuronal function, we inhibited the function of the *TUNAR*-PE1 (Figure 5c) and assessed the effect on neuronal firing. We found that when the activity of this *TUNAR* enhancer is inhibited, there is an increase in burst duration (Figure 6c). In addition, as the neurons mature, we detected an overall increase in the weighted mean firing rate of the neurons when *TUNAR*-PE1 is inhibited (Figure 6b). This may be due to an increase in burst duration combined with an increase in the firing rate within bursts (Figure S7i). We conclude that inhibiting the function of a single human-biased activity-dependent enhancer affects the firing properties of neurons.

## Discussion

We used cultured human-chimpanzee tetraploid neurons to identify activity-dependent genes that are differentially induced in human compared to chimpanzee piNs within the same nuclear environment. Neuronal activity drives the maturation of cortical excitatory neurons, a process that is dramatically slowed and more complex in humans compared to chimpanzees and other species. We employed a highly reproducible excitatory neuron differentiation protocol to generate relatively homogeneous cell populations from diploid and tetraploid iPSCs and delivered a strong, synchronous stimulus to neurons by depolarizing the membrane with elevated levels of KCl. Both the homogeneity of the culture and the consistency of the differentiation protocol across diploid and tetraploid cells allowed us to make comparisons between species that have not previously been possible (Agoglia et al., 2021). This allowed us, for the first time, to make a comprehensive and in-depth mechanistic comparison between ADGE programs of human and chimpanzee neurons, and identify genes for which sequence changes in nearby regulatory regions drive these evolutionary changes.

We identified over 200 genes that are activity-regulated and show significant species bias. A subset of these genes are activity-dependent only in one species, which may be due to the gain or loss of stimulus-responsive regulatory elements. Other genes are activity-dependent in both species, but show consistently higher expression in one throughout the time course, suggesting an alternate mechanism where evolution has acted on a constitutive enhancer. Notably, we found that a larger number of LRGs show species-specific changes than IEGs. Between human and chimpanzee neurons, we detected no significant difference in the expression levels of any canonical IEG TFs, including AP-1 and NR4A family members, suggesting that evolution has tinkered with the expression of downstream effector genes, but not the master regulators of the membrane depolarization response. Prior studies have shown that IEG programs are highly similar across cell types, while LRG programs are highly cell type specific (Hrvatin et al., 2018; Traunmüller et al., 2025). We have extended this finding beyond differences between cell types in the same species to differences between species in the same cell type. Given the cell type specificity of LRG expression, it will be worthwhile to uncover human-specific features of ADGE in additional neuronal subtypes differentiated from tetraploid cells.

Our finding that the majority of species-biased and activity-dependent open chromatin regions contained AP-1 motifs, and that the majority of FOS binding sites show a species bias in binding suggests that the evolution of ADGE occurs largely through changes in FOS binding sites. This includes a similar number of gained and lost binding sites in human piNs, suggesting that changes in FOS binding sites may direct the downstream expression of species-specific activity-dependent genes across species more generally. This is in contrast to recent work that found that AP-1 motifs are enriched in human-specific but not chimpanzee-specific open chromatin regions in human brain tissue (Caglayan et al., 2023). Our work shows that FOS binding, and not just AP-1 accessibility, changes through evolution and alters downstream activity-dependent gene expression.

Through the study of human-chimpanzee tetraploid neurons, we were able to identify gene expression changes between species that are due to nearby *cis* changes in gene regulatory elements. The AP-1 motif is highly constrained in human populations, and a single base change at six out of the seven motif positions leads to a strong reduction in the ability of FOS to bind (Yang et al., 2022). We found that many species-biased FOS binding sites have single base changes in AP-1 motifs, and we demonstrated that recreating those evolutionary sequence changes in a reporter assay dramatically affects enhancer activity. Thus, AP-1 sites appear to be abundant targets of evolutionary changes in activity-dependent enhancer activity and can gain or lose activity through single base pair changes in evolution.

Testing the function of enhancers in their endogenous context has historically been challenging, given the limited tools for silencing enhancers and the difficulty in linking putative enhancers to target genes. Here we searched for enhancer-gene pairs within the limited space of neuronal activity-dependent genes and regulatory regions, which made linking species-biased activity-dependent genes and FOS binding sites somewhat simpler. We were able to show using CRISPRi that targeting FOS binding sites at enhancers near *TUNAR* and *PCSK1* significantly reduce activity-dependent expression of these genes, thus implicating these specific enhancers in control of nearby genes.

In addition to *cis*-regulatory changes, we identified a number of *trans*-regulated genes and open chromatin regions in piNs. These *trans*-regulated genes and sites may be regulated by TFs with species-biased expression as described in (Song et al., 2025), or by TFs that have species-specific amino acid changes. *Trans*-regulated and activity-dependent open chromatin regions are also enriched for the AP-1 motif. Notably, there is a human-derived SNV in the FOS coding sequence that leads to a nonsynonymous amino acid change in the protein compared to chimpanzees. Thus, it is of interest in the future to test what effect FOS may have on regulating genes in *trans*, through differences in the abundance of its binding partners or perhaps through changes in its amino acid sequence.

We hypothesized that evolutionary changes in the expression of activity-dependent genes may modify activity-dependent responses in neurons, changing synaptic responses and ultimately contributing to the slowed pace of maturation in human neurons. The species-biased genes that we identified have myriad functions, suggesting that many synaptic programs may have been affected by changes in the ADGE response during human evolution. This is supported by our MEA results that found that both *TUNAR* and *SV2C* knockdown led to changes in the length of firing bursts across neuron maturation. Combined with their known roles in regulating synaptic signaling and calcium dynamics (Schivell et al., 2005; Dardou et al., 2013; Dunn et al., 2017; Senís et al., 2021), this suggests that human-biased ADGs may dampen calcium signaling, whether cell-autonomously through pTUNAR-mediated calcium sequestration or by limiting the strength of synaptic inputs via an SV2C-dependent block of vesicle release. Interestingly, calcium signaling has long been known to regulate neuronal maturation, and recently published studies have implicated the regulation of calcium influx in driving the prolonged period of human neuron maturation and axonal outgrowth (Ciceri et al., 2024; Hergenreder et al., 2024; Lindhout et al., 2025). Future studies in the mouse or in human brain organoids should further elucidate the mechanisms by which these genes modulate neuronal function.

Finally, by targeting *TUNAR*’s human-biased activity-dependent enhancer with CRISPRi, we showed that modification of a single FOS binding site in human neurons can affect the firing properties and perhaps the maturation rate of the neurons. This further underscores the role that the evolution of FOS binding sites plays in the modification of activity-dependent gene programs between species, and suggests that just a small number of species-specific sequence changes in an enhancer can have measurable effects on neuronal function.

Key driving forces in evolution are a changing environment and an organism’s ability to respond to external cues from that environment. Neurons respond to environmental stimuli in the short term by communicating with one another at synapses, and in the long term, by activating ADGE, in part via FOS-mediated enhancer activation. While our study focused on the evolution of human-biased ADG programs in excitatory neurons, AP-1 factors play crucial roles across a variety of cell types. AP-1 mediates stimulus-responses throughout the brain and in many different neuronal subtypes, as well as in the immune system, pancreatic beta cells, and fibroblasts (Greenberg and Ziff, 1984; Glauser and Schlegel, 2007; Wagner and Eferl, 2005; Yap and Greenberg, 2018; Hrvatin et al., 2018; Atsaves et al., 2019). Our results point to a possible interaction between cell-type-specific TF binding and AP-1 site evolution. AP-1 site evolution may therefore be a widespread mechanism for the evolution of stimulus responses at the cellular level. A recent and growing body of work also suggests that AP-1 may play a role in dictating the differentiation, maturation, and senescence of cells across the body (Martínez-Zamudio et al., 2020; Byrns et al., 2021; Herring et al., 2022; Patrick et al., 2024; Ciceri et al., 2024). These findings suggest that the evolution of AP-1 sites may concurrently affect the cell’s stimulus response repertoire and ultimately an organism’s life history. As differentiation protocols expand, the tetraploid stem cell model can serve as a critical tool for further exploring these possibilities.

## Acknowledgments

The authors thank all members of the M.E.G. and C.A.W. labs for helpful insight. The authors thank the Boston Children’s Hospital Human Neuron Core, the Harvard Medical School Department of Immunology FACS core, the Harvard Medical School Core for Imaging Technology and Education, and the Harvard Medical School Neurobiology Imaging Facility. We thank Dr. Wade Harper at Harvard Medical School for generously sharing plasmids to make stable NGN2-expressing cell lines, and Dr. Martin Kampmann for sharing CRISPRi cell lines. C.A.W. and M.E.G. are supported by the Allen Discovery Center program, a Paul G. Allen Frontiers Group advised program of the Paul G. Allen Family Foundation, and the Hock E. Tan and K. Lisa Yang Center for Autism Research at Harvard University. A.C.C. is supported by the Hanna H. Gray Fellowship from the Howard Hughes Medical Institute. J.H.T.S. is supported by the Y. Eva Tan Fellowship from the Tan Yang Autism Center at Harvard University, the Howard Hughes Medical Institute Fellowship from the Helen Hay Whitney Foundation, and grant no. K99MH136290 from NIMH/NIH. C.A.W. is supported by the Templeton Foundation. C.A.W. and D.M.K. are Investigators of the Howard Hughes Medical Institute.

## Author Contributions

**Ava C. Carter**: Conceptualization, Methodology, Formal Analysis, Investigation, Writing - Original Draft, Writing - Review & Editing, Visualization. **Gabriel T. Koreman**: Conceptualization, Methodology, Formal Analysis, Investigation, Writing - Original Draft, Writing - Review & Editing, Visualization. **Jillian E. Petrocelli**: Investigation. **Josephine E. Robb**: Investigation. **Evan M. Bushinsky**: Investigation. **Sara K. Trowbridge**: Investigation. **David M. Kingsley**: Resources, Writing - Review & Editing. **Christopher A. Walsh**: Conceptualization, Writing - Original Draft, Writing - Review & Editing. **Janet H.T. Song**: Conceptualization, Methodology, Formal Analysis, Investigation, Writing - Original Draft, Writing - Review & Editing, Visualization. **Michael E. Greenberg**: Conceptualization, Writing - Original Draft, Writing - Review & Editing.

## Materials and Methods

### PSC culture

Diploid cell lines were generated in the lab of Yoav Gilad. NCRM5 NGN2 dCas9-KRAB cell line was a gift from the lab of Martin Kampmann (Tian et al., 2019). They were cultured under the ESCRO protocol E00041. iPSCs were cultured on hESC-qualified Matrigel (Corning # 354277) in mTeSr Plus (STEMCELL Technologies # 100-0276). They were passaged using Dispase (Life Technologies # 17105041) or Accutase (Thermo Fisher Scientific # A1110501) as needed every 3-4 days and supplemented for one day with 10µM ROCK-inhibitor Y-27632 (STEMCELL Technologies # 72304). The media was changed every 1-2 days.

### Engineering NGN2 cell lines

To make NGN2 lines from human diploid, chimpanzee diploid, human autotetraploid, chimpanzee autotetraploid, and allotetraploid iPSCs (Gallego Romero et al., 2015; Song et al., 2021), we inserted a doxycycline-inducible NGN2 cassette into the AAVS1 locus (Tian et al., 2019) as described in (Song et al., 2025). Briefly we in vitro transcribed an sgRNA targeting the AAVS2 locus (5’-GGGGCCACTAGGGACAGGAT-3’) and complexed it with Cas9-NLS protein (QB3, UC Berkeley). We delivered the complexed RNP with a donor plasmid pAAVS1-TRE3G-NGN2 (Ordureau et al., 2020) (gift from Wade Harper Lab) into 1 million iPSCs with the Neon Transfection Kit (Thermo Fisher # MPK10096). Cells were plated in mTeSR Plus (STEMCELL Technologies # 100-0276) with 1 µM thiazovivin (Tocris # 3845) in 10cm plates for 2-3 days. Cells containing the donor plasmid were then selected using 0.5mg/mL puromycin (Thermo Fisher # 10131035) for 3-5 days and then allowed to recover for 7-10 days without puromycin. Individual colonies were picked into a 96-well plate using the EVOS M5000 microscope (Thermo Fisher) and genotyped with primers spanning the 3’ and 5’ junctions (Table S1). Clones that were positive by PCR were then Sanger sequenced to confirm the genotype and karyotyped by WiCell.

### Generation of sgRNA-expressing cell lines

sgRNA sequences targeting genes of interest were gathered from Horlbeck et al., 2016 (Horlbeck et al., 2016) or generated using CRISPick (Doench et al., 2016; Sanson et al., 2018) if not present in the Horlbeck data set. sgRNA sequences can be found in Table S1. sgRNAs were ordered as two oligos, with 5’-TTG and GTTTAAGAGC-3’ overhangs on the top oligo and 5’-TTAGCTCTAAAC and CAACAAG-3’overhangs on the bottom oligos. Oligo pairs were phosphorylated and annealed with T4 PNK for 30 minutes at 37 ^°^C followed by 5 minutes at 95 ^°^C and then a ramp down to 25 ^°^C at a rate of 1 ^°^C/s. pBA904 (Replogle et al., 2020) (Addgene # 122238) was digested first with BlpI (New England Biolabs # R0585S) in CutSmart Buffer (New England Biolabs # B7204S) and then by BstXI (New England Biolabs # R0113S) in NEBuffer r3.1 (New England Biolabs # B6003S) and cleaned up with gel extraction. Annealed oligos were diluted 1:200 and then ligated into pBA904 using T4 DNA ligase (New England Biolabs # M0202T). Ligated vectors were transformed and grown in STBL3 cells and then purified and sequenced to confirm the correct sgRNA sequence was inserted using sanger sequencing or whole plasmid sequencing.

To make stable sgRNA-expressing cell lines, we first made lentivirus from pBA904 containing our sgRNA sequences. Lentivirus was packaged in LentiX 293T cells (Takara Bio # 632180) using VSVG and PSPAX2 following calcium phosphate transfection. Lentivirus-containing media was harvested 48 hours after transfection and concentrated using 40% PEG-8000 (Sigma Aldrich # P5413-2KG) and 1.2M NaCl in PBS. Lentivirus was added after one freeze-thaw cycle to human or chimpanzee PSCs that constitutively express dCas9-KRAB from the CLYBL locus. After 72 hours, infected cells were disscociated using Accutase (Thermo Fisher Scientific # A1110501) and FACS sorted for cells expressing blue fluorescent protein (BFP) from pBA904. BFP+ cells containing each sgRNA construct were re-plated and cultured as stable PSC lines.

### piN differentiation

piNs were differentiated using a protocol modified from (Nehme et al., 2018). iPSCs were dissociated on day 0 using Accutase (Thermo Fisher Scientific # A1110501) and plated at a density of 50,000 cells/cm^2^ for diploids and 25,000 cells/cm^2^ for tetraploids on tissue culture dishes coated with growth factor-reduced Matrigel (VWR # 47743-720) in mTesr+ media (STEMCELL Technologies # 100-0276) supplemented with 10µM ROCK-inhibitor Y-27632 (STEMCELL Technologies # 72304). On day 1, the media was changed to KSR (Knockout DMEM [Thermo Fisher Scientific # 10829018], with 15% Knockout Serum Replacement [Life Technologies # 10828028], 1% L-Glutamine [Life Technologies # 25030081], 1% MEM-NEAA [Life Technologies # 11140050], 1% Penicillin/Streptomycin [Life Technologies # 15140122], 0.1% beta-mercaptoethanol [Life Technologies # 21985023]) with 2 µg/mL doxycycline (Sigma # D9891), 10µM SB431542 (Sigma # S4317), 2µM XAV939 (STEMCELL Technologies # 72672), and 0.1µM LDN193189 (STEMCELL Technologies # 72147). On day 2, the media was changed to half KSR, half NIM (DMEM/F-12 [Life Technologies # 11320082], 1x N2 Supplement-B [STEMCELL Technologies # 07156], 1% MEM-NEAA, 1% Penicillin/Streptomycin, 1% GlutaMAX [Life Technologies # 35050061], 0.16% D-Glucose), supplemented as on day 1 with doxycycline, SB, XAV, and LDN. On day 4, the cells were dissociated with Accutase and replated at a density of 40,000 cells/cm^2^ for diploids and 20,000 cells/cm^2^ for tetraploids on growth-factor reduced Matrigel in complete neurobasal media (Neurobasal [Thermo Fisher Scientific # 21103049], 1% MEM-NEAA, 1% Penicillin/Streptomycin, 1% GlutaMAX [Life Technologies # 35050061], 1x N2 Supplement-B [STEMCELL Technologies # 07156], 1x B27 without vitamin A [Life Technologies # 12587010], 0.1% Mouse laminin [Life Technologies # 23017015], 20ng/mL rhBDNF [Peprotech # 450-02], 10ng/mL rhGDNF [Peprotech # 450-10], 1µM L-Ascorbic Acid [Sigma # A4544], 20µM Dibutyryl cAMP [Sigma # D0260]) supplemented with 10µM ROCKi and 2µg/mL doxycycline. cNB Media + 2µg/mL doxycyline was then changed once per week (Days 5, 12, 19, 26) and 2µg/mL doxycycline was spiked into the media halfway through the week (Days 8, 15, 22). For RNA-seq, ATAC-seq, and CUT&Tag, 0.05µM AraC (Sigma # C6645) was added from day 12-14 to remove any remaining undifferentiated mitotic cells.

### KCl stimulation of piNs

All KCl stimulation experiments were performed on day 28 of differentiation unless otherwise noted. On day 27, piNs were silenced with 1 µM TTX (VWR # 102663-330) and 100 µM APV (Thermo Fisher Scientific # 010610) to reduce background activity for 16-20 hours. On day 28, 170 mM KCl depolarization buffer (170 mM KCl, 2 mM CaCl_2_, 1 mM MgCl_2_, 10 mM HEPES) was added to a final concentration of 55 mM. For experiments in which piNs were depolarized for different amounts of time, KCl was added at different times during the day so that collection and downstream assays could be performed at one time for all samples.

### qRT-PCR

For RNA collection, cell culture wells were washed with 1X PBS (GIBCO # 10010023), and then cells were scraped in 0.5-1mL TRIzol (Life Technologies # 15596026) and frozen at −80 ^°^C. After thawing 0.2 volumes of chloroform were added and sample was homogenized. RNA was then harvested from the aqueous phase using the Qiagen RNEasy Micro kit. 250-500ng of RNA was then reverse transcribed into cDNA using the SuperScript VILO kit (Life Technologies # 11755050). qPCR was performed on an Applied Biosystems QuantStudio 3 Real-Time PCR System using PowerUP SYBR Green Master Mix (Fisher Scientific # A25778). Primer sequences can be found in Table S1.

### RNA-seq

For RNA collection, cell culture wells with diploid, autotetraploid, and allotetraploid piNs were washed with 1X PBS, and then cells were scraped in 0.5-1mL TRIzol (Life Technologies # 15596026) and frozen at −80 ^°^C. After thawing 0.2 volumes of chloroform were added and sample was homogenized. RNA was then harvested from the aqueous phase using the Qiagen RNEasy Micro kit. 10ng RNA was used as input to the SMARTer Stranded Total RNA-Seq Kit v2 - Pico Input Mammalian kit (Takara Bio # 634413). RNA-seq libraries were quantified using the Qubit DS DNA high-sensitivity assay kit (Fisher Scientific # Q32854) and sequenced with an Illumina NovaSeq 6000 with paired-end 150bp reads. A table with sample numbers and associated metadata is found in Table S2.

For RNA-seq of *TUNAR* and *SV2C* CRISPRi piNs, stable iPSC lines generated from two human iPSC lines (H2-C and NCRM5) containing nontargeting sgRNAs (2 NT sgRNAs), sgRNAs targeting the *TUNAR* TSS (2 *TUNAR* TSS sgRNAs) or the *SV2C* TSS (1 pool of 3 sgRNAs targeting the TSS) were differentiated into piNs for 28 days and RNA was harvested in TRIzol. Sequencing libraries were prepared as described above. Samples were sequenced with an Illumina NextSeq 500 with paired-end 37bp reads. A table with sample numbers and associated metadata is found in Table S5.

### ATAC-seq

ATAC-seq was performed as described in (Corces et al., 2017) with minor modifications. For ATAC-seq, 250µL lysis buffer (10mM Tris-HCl pH7.4, 10mM NaCl, 3mM MgCl_2_, 0.1% NP-40, 0.1% Tween-20) was added to each 12-well well of piNs at D28. Cells were scraped and collected into conical tubes in a final volume of 500µL lysis buffer and then incubated at 4 ^°^C rotating to release nuclei, followed by centrifugation at 830 rcf for 10 minutes. Nuclei were then resuspended in 1mL of lysis buffer and counted. 50,000 nuclei per sample were aliquoted and centrifuged for 10 minutes at 830 rcf. After removing the supernatant, the pellet was resuspended in transposition mix (1X TD buffer [10mM Tris-HCl pH7.6, 5mM MgCl_2_, 10% dimethyl formamide], 0.33x PBS, 0.001% Digitonin, 0.1% Tween-20, 100nM Tn5 transposase [made in house]), and incubated for 30 minutes at 37 ^°^C. After transposition, DNA was immediately purified using the Zymo DNA Clean and Concentrator Kit (Research Products International # ZD4014) and eluted in 21 µL water.

### ATAC-seq library preparation and sequencing

For ATAC-seq library preparation, the entire 20µL eluate was added to a PCR reaction consisting of 1x NEBNext High Fidelity PCR Master Mix (New England Biolabs # M0541L) 1.25µM Ad1 and 1.25µM Ad2 (Table S1), and amplified for 8 cycles (72 ^°^C for 5 min, 98 ^°^C for 30 seconds followed by 5 cycles of 98 ^°^C for 10 seconds, 63 ^°^C for 30 seconds and 72 ^°^C for 1 minute). Libraries were purified using the Zymo DNA Clean and Concentrator Kit (Research Products International # ZD4014), quantified with a KAPA Library Quantification Kit (Roche Diagnostics # 07960140001) and sequenced with an Illumina NovaSeq 6000 with paired-end 150bp reads. A table with sample numbers and associated metadata is found in Table S3.

### CUT&Tag

For CUT&Tag, tetraploid piNs that were unstimulated or stimulated for 2hr with 55mM KCl were scraped off of tissue culture plates on D28 in NE1 lysis buffer (20mM HEPES pH 7.9, 10mM KCl, 0.1% Triton X-100, 3mM MgCl_2_, 0.5mM Spermidine, 1 tablet Complete Protease Inhibitor EDTA-free per 50mL). Nuclei were isolated by rotating for 10 minutes at 4 ^°^C followed by centrifugation at 200 rpm at 4 ^°^C in a swinging bucket centrifuge. Nuclei were resuspended in 1mL WB150 (20mM HEPES pH 7.5, 150mM NaCl, 0.2% Tween-20, 1g/L BSA, 0.5mM Spermidine [Sigma Aldrich # S2501], 10mM sodium butyrate [Sigma Aldrich # 19-137], 1 tablet Complete Protease Inhibitor EDTA-free [Roche Diagnostics # 11873580001] per 50mL) and counted on a hemocytometer. 500,000 nuclei per CUT&Tag were aliquoted and the volume of WB150 was brought up to 1mL per reaction. 20µL Concanavalin A bead (Bangs Laboratories # BP531) slurry was prepared for each CUT&Tag reaction by washing twice in binding buffer (20µM HEPES-KOH pH 7.9, 10µM KCl, 1µM CaCl_2_, 1µM MnCl_2_), and then resuspending in 10µL binding buffer per reaction. 10µL beads were added to the nuclei and incubated for 10 minutes at room temperature. The supernatant was then removed using a magnet, and 50µL antibody buffer (WB150 supplemented with 0.1% Triton X-100 and 2mM EDTA) containing 1µL primary antibody (FOS: Cedarlane # 226003; H3K27Ac: Abcam # ab4729; IgG: Cell Signaling Technology # 2729S) was added. Samples were incubated overnight at 4 ^°^C. On day two, buffer containing primary antibody was removed and 100µL of antibody buffer containing 1.2µL anti-rabbit (Fisher Scientific # NBP172763) or anti-mouse (Abcam # ab46540) secondary antibody was added with trituration to resuspend the beads. After 1 hour at room temperature, beads were washed 3 times with 1mL WB150 and then resuspended in 100µL WB300 (20mM HEPES pH 7.5, 300mM NaCl, 0.2% Tween-20, 1g/L BSA, 0.5mM Spermidine, 10mM sodium butyrate, 1 tablet Complete Protease Inhibitor EDTA-free per 50mL) containing 2.5µL proteinAG-Tn5 (Epicypher Inc # SKU: 15-1017). After incubating for 1 hour at room temperature, beads were washed 3 times with 1mL WB300 and then resuspended in tagmentation buffer (WB300 supplemented with 10mM MgCl_2_). Samples were incubated for 1 hour at 37 ^°^C, and then tagmentation was stopped by adding 17mM EDTA, 0.1% SDS, and 0.17mg/mL proteinase K (Life Technologies # AM2546) followed by incubation for 1 hour at 50 ^°^C shaking at 500 rpm. DNA was then extracted twice with Phenol-Chloroform, precipitated twice with ethanol and 0.02mg/mL GlycoBlue (Life Technologies # AM9515), and re-suspended in 22µL of water.

### CUT&Tag library preparation and sequencing

For CUT&Tag library preparation, 20µL of precipitated DNA was added to 13.5µL water, 10µL 5x KAPA HiFi Buffer (Roche Diagnostics # KK2101), 1.5µL 10mM dNTPs (Roche Diagnostics # KK2101), 1µL KAPA HiFi Polymerase (Roche Diagnostics # KK2101), 2µL 10µM Ad1, and 2µL 10 µM indexed Ad2 (Table S1). The reaction was incubated at 72 ^°^C for 5 minutes, then 98 ^°^C for 30 seconds, followed by 14 cycles of 98 ^°^C for 10 seconds, 63 ^°^C for 30 seconds, and 72 ^°^C for 1 minute. Samples were then incubated at 72 ^°^C for 5 minutes. Following PCR, samples were purified using double-sided SPRI beads (Ampure XP, Thermo Fisher Scientific # NC0110018). 25µL of SPRI beads were added to the PCR reaction and incubated at room temperature for 8 minutes followed by capture on a magnet for 8 minutes. Supernatant was then recovered and moved to a fresh tube. 65µL SPRI beads were then added to the supernatant and incubated at room temperature for 8 minutes followed by capture on a magnet for 8 minutes. Supernatant was then discarded and beads were washed twice for 30 seconds each with 200µL 80% ethanol, and then dried. Beads were then resuspended in 53µL 10mM Tris-HCl pH 7.5 and then captured on a magnet. 50µL supernatant was moved to a new tube for the final SPRI cleanup. 55µL SPRI beads were added to the supernatant, incubated and then washed twice as described above with 80% ethanol. DNA was eluted in 20µL of 10mM Tris-HCl pH 7.5. Libraries were quantified with the KAPA Library Quantification kit (Roche Diagnostics # 07960140001) and sequenced with an Illumina NovaSeq 6000 with paired-end 150bp reads. A table with sample numbers and associated metadata is found in Table S4.

### Immunofluorescence

After 28 days of differentiation on glass coverslips, piNs were fixed with 4% PFA/4% sucrose solution for ten minutes at room temperature. After fixing, cells were washed 3x with 1x PBS and blocked for 1 hour in 0.05% donkey serum (Sigma Aldrich # D9663) and 0.2% Triton X-100 in PBS at room temperature. Cells were incubated in primary antibody (MAP2: Lifespan Biosciences # LS-C61805) in 0.05% donkey serum overnight at 4 ^°^C. After washing 3 times in PBS with 0.1% Triton X-100, cells were incubated in secondary antibody (anti-chicken Invitrogen # A-11039) in 0.05% donkey serum for 1 hour at room temperature. After washing 3 times in PBS, samples were mounted on slides using DAPI fluoromount G (Thermo Fisher Scientific # OB010020), sealed with clear nail polish, and imaged on a Leica Sp8 confocal microscope.

### Multielectrode array

A 48-well CytoView MEA plate (Axion Biosystems # M768-tMEA-48B) was coated with 150µL of 0.01% PEI (Sigma Aldrich # 408727) in Borate Buffer (Life Technologies # 28341) for one hr, washed 4x with sterile water and allowed to dry, covered overnight in the hood. The following day 150µL of 0.01mg/mL mouse laminin (Life Technologies # 23017015) was added to each well in DPBS with Ca2+/Mg2+ (Life Technologies # 14040133) and incubated overnight at 37 ^°^C. On Day 4 of the neuron differentiation, Human iPSC derived astrocytes (Ncardia # K0101) were thawed and 9000 were immediately plated in 150µL of complete neurobasal (Neurobasal [Thermo Fisher Scientific # 21103049], 1% MEM-NEAA, 1% Penicillin/Streptomycin, 1% GlutaMAX [Life Technologies # 35050061], 1x N2 Supplement-B [STEMCELL Technologies # 07156], 1x B27 without vitamin A [Life Technologies # 12587010], 0.1% Mouse laminin [Life Technologies # 23017015], 20ng/mL rhBDNF [Peprotech # 450-02], 10ng/mL rhGDNF [Peprotech # 450-10], 1µM L-Ascorbic Acid [Sigma # A4544], 20µM Dibutyrul cAMP [Sigma # D0260]) supplemented with 10µM ROCKi and 2µg/mL doxycycline) in each well. piNs were cultured as described up to replating on Day 4. On Day 5, neurons were treated with 60 nM AraC (Sigma # C6645) for two days. On Day 7, neurons were lifted from plates using Accutase and mixed with a newly thawed vial of passaged Human iPSC derived astrocytes (Ncardia # K0101) at a ratio of 9000 astrocytes to 60,000 neurons. This mixture was then spot-plated in 20µL of complete neurobasal supplemented with 10µM ROCK inhibitor (STEMCELL Technologies # 72304) in each well of the 48-well MEA plate described above. Cells were allowed to adhere for one hour at 37 ^°^C before an additional 150µL of cNB supplemented with 10µM ROCK inhibitor was added to each well. The following day, 150µL of cNB was added to each well. Half media changes were performed every 3 days with cNB until around Day 14, after which media was replaced with Complete Brainphys media (BrainPhys Media [STEMCELL Technologies # 05796], 1% Penicillin/Streptomycin, 1x N2 Supplement-A [STEMCELL Technologies # 07152], 1x NeuroCult™ SM1 Without Vitamin A [STEMCELL Technologies # 05731], 0.1% Mouse laminin [Life Technologies # 23017015], 20ng/mL rhBDNF [Peprotech 450-02], 10ng/mL rhGDNF [Peprotech # 450-10], 1µM L-Ascorbic Acid [Sigma # A4544], 20µM Dibutyrul cAMP [Sigma # D0260], 2ug/mL doxycycline [Sigma # D9891]). Half media changes were completed almost every 3 days, immediately following recording.

Recordings were completed using the Axion Maestro Pro (Maestro-625) recording system. Plates were first allowed to equilibrate to 37 ^°^C and 5% CO2 after transfer for 10 minutes and recordings were done immediately for 10 min. Voltage activity was recorded at 12.5kHz and passed through a 3kHz Kaiser-Window low pass filter and 200Hz IIR high pass filter. Spikes were counted using an adaptive threshold crossing model with a standard deviation threshold of 6. Bursts were detected using a maximum inter-spike interval (ISI) method threshold with a maximum ISI of 100ms and a minimum of 5 spikes. Active electrodes were those that had at least 5 spikes per minute.

### Multielectrode array analysis

CSV files containing recording metadata and metrics were made using the AXIS Neural Metric tool. These files were analyzed in R to determine metric evolution across recordings. Wells were excluded if they had less than 10 active electrodes or less than 15 covered electrodes. Wells were grouped depending on their expressed sgRNA (non-targeting guides 1 and 2 were combined, as were *TUNAR* TSS guides 1 and 2, and *TUNAR*-PE1 guide 4 and guide 5). Differential metrics were identified by performing ANOVAs between linear mixed effect models with and without a sgRNA fixed effect. All models included a nested, plate:well random effect and day count fixed effect. Significant metrics (p < 0.05, Holm-Sidak corrected) were chosen and modeled with a sgRNA fixed effect. Estimated marginal means (emmeans) were found for each sgRNA group and compared pairwise across days using the emmeans ‘pair-wise’ function in R (Holm-Sidak corrected p-values reported).

### Mouse cortical neuron dissection and culture

Mouse cortical neurons were prepared as previously described (Pollina et al., 2023) and all animal use was approved and overseen by the Harvard University Institutional Animal Care and Use Committee. Briefly, cortices were dissected from approximately E16.5, mixed-sex embryos (C57/BL6) with each biological replicate coming from a different dam and dissociation procedure. Cortices were digested using Papain (Sigma Aldrich # 10108014001) in the presence of DNAse (Sigma Aldrich # DN25-10MG), digestion was stopped through addition of ovomucoid (Worthington Biochemical Corp. # LS003085), and cortices were triturated using a P1000 pipette tip. Neurons were plated on precoated (20 µg/mL Poly-d-lysine (Sigma Aldrich # P7405-5MG), 4 µg/mL Mouse Laminin (Life Technologies # 23017015) 24-well tissue culture treated plates and subsequently cultured in Neurobasal [Thermo Fisher Scientific # 21103049] supplemented with 1% Penicillin/Streptomycin [Life Technologies # 15140122], 1% GlutaMAX [Life Technologies # 35050061], and 1x B27 without vitamin A [Life Technologies # 12587010] at a density of 150k cells/cm ^2^. Half the media was changed at DIV 3.

### Luciferase construct cloning

Luciferase constructs are all based on the NanoLuc® luciferase reporter vector pNL3.2[NlucP/minP] (Promega # N104A). Enhancers were amplified from human and chimpanzee genomic DNA, or synthesized as gene blocks and cloned in directly upstream of the minimal promoter via PCR amplification (when using gene blocks) or HindIII digestion (New England Biolabs # R3104T) (when not using gBlocks) of pNL3.2 followed by In-Fusion Snap Assembly (Takara Bio # 638949). All PCR reactions used NEBNext High-Fidelity 2X PCR Master Mix (New England Biolabs # M0541L). Gene block and oligo sequences can be found in Table S1. Site-directed mutagenesis was performed using the QuikChange Lightning Site-Directed Mutagenesis Kit (Agilent # 210518) on the existing nano-luciferase plasmids containing the HBF-226 and HBF-1033 enhancers. Oligo sequences can be found in Table S1.

### Luciferase assays

DIV6 neurons were transfected with 500ng of enhancer-containing nanoluciferase plasmid, 20ng of pBluescript II SK+ (Stratagene # 212205) for added DNA load, 70ng of an EGFP expression vector for visualization of transfection efficiency (FUGW, Addgene # 14883), and 10ng of pGL4.53[luc2/PGK] (Promega # E5011) as a normalizing luciferase control. The empty pNL3.2[NlucP/minP] (Promega # N1041) construct was used as a negative control while a highly-inducible enhancer (Enhancer 39, (Malik et al., 2014)) was used as a positive control. Neurons were transfected for two hours using Lipofectamine 2000 (Life Technologies # 11668500) in non-supplemented Neurobasal media (Thermo Fisher Scientific # 21103049) and were returned to a 1:1 mixture of fresh and conditioned, supplemented Neurobasal media (Neurobasal [Thermo Fisher Scientific # 21103049] supplemented with 1% Penicillin/Streptomycin [Life Technologies # 15140122], 1% GlutaMAX [Life Technologies # 35050061], and 1x B27 without vitamin A [Life Technologies # 12587010]). Neurons were silenced on DIV7 with1µM TTX (VWR # 102663-330) and 100µM APV (Thermo Fisher Scientific # 010610) to reduce background spontaneous activity for 16 hours. DIV8 neurons were stimulated through the addition of 170mM KCl depolarization buffer (170 mM KCl, 2 mM CaCl_2_, 1 mM MgCl_2_, 10 mM HEPES) to a final concentration of 55mM for 6hr. Following stimulation, neurons were collected through the addition of 110µL of Passive Lysis Buffer (Promega E1941), shaking at 225 RPM for approximately one hour, and scraped from the wells. Two 24-well wells were used for each timepoint (no stimulation, 6hr KCl stimulation) and collected separately. Following collection, lysates were stored at − 20 ^°^C for up to one week before thawing, spinning down for two minutes, and performing the Nano-Glo Dual-Luciferase Reporter Assay System (Promega # N1620) according to the manufacturer’s supplied documentation. The lysate from each 24-well of the original culture was run in a separate well for measuring luminescence. Luminescence was measured using a BioTek Synergy H1 plate reader. Nano-luciferase luminescence was first normalized to control firefly luciferase luminescence to obtain a normalized ratio for each well. This ratio was then normalized to the negative control (empty vector) ratio for both conditions. Assays where the positive control’s double-normalized ratio did not show greater than a 10-fold increase upon KCl stimulation (indicative of poor stimulation), were discarded from future analysis.

### RNA-seq Analysis in diploid, autotetraploid, and allotetraploid piNs

RNA-seq from diploid, autotetraploid, and allotetraploid piNs was analyzed as previously described (Song et al., 2021). Trimgalore v0.6.6 and cutadapt v2.5 (Martin, 2011) were used to trim adapter sequences and fastqc v0.11.5 was used to assess sequencing quality. Star v2.7.9a (Dobin et al., 2013) with two-pass mapping was used to align reads to a composite human-chimpanzee genome (from hg38 and panTro6) and the number of reads that mapped uniquely (MAPQ=255) to each gene was counted with featureCounts from subread v2.0.0 (Liao et al., 2014).

The following piN lines were removed from downstream analyses due to a failure to induce FOS expression with 2hr KCl treatment: C3-C, H1H1a_H6-1, C1_A4-3, C2_C8-1, H1H1a_D3-2.

Gene annotations for featureCounts (“byexon-gene” and “bysnp-gene”) were previously generated in (Song et al., 2021), and were further filtered as follows. Exons or SNVs where 10% or more of reads from diploid and autotetraploid lines map to the incorrect species in piNs were removed. In addition, we filtered out genes where 10% or more of the reads mapped incorrectly in piNs when quantifying counts from exons and SNVs separately. We also combined exons and SNVs into a single annotation where only SNVs that did not overlap exons were included. We removed genes for which more than 10% of reads from diploid and autotetraploid piNs mapped to the incorrect species. The resulting annotations contained 477,300 exons and 73,993 SNVs from 98,684 genes. After alignment, we performed differential expression (DE) analysis and allele-specific expression analysis (ASE) using DESeq2 (Love et al., 2014) with default parameters. Genes were considered significantly differential if the p-adjusted after Benjamini-Hochberg FDR correction was p<0.05. We classified genes by regulatory type as follows: *cis*: significant ASE, significant DE, Log_2_(FC) in the same direction between ASE and DE. *trans*: significant DE, not significant ASE. *compensatory*: significant ASE, not significant DE.

We adjusted read counts by feature length to account for any differences between length in the chimpanzee and human genomes. This produced nearly identical results to those without length correction. Only genes that we found to be significantly DE or ASE with and without length correction were kept, but visualization was performed without length correction. We used a cutoff of FC > 1.5 for all downstream analyses of gene sets. For activity-dependent gene calling, we used the same parameters to call DE between unstimulated and stimulated samples. Associated data can be found in Table S2.

### Analysis of tetraploid-biased gene expression

To assess whether tetraploidization affects gene expression in piNs, we compared human diploid to human autotetraploid neurons and chimpanzee diploid to chimpanzee autotetraploid neurons. Although there are very few differences in gene expression between diploid and autotetraploid iPSCs (Song et al., 2021) or NPCs (Song et al., 2025), we found that there are 2434 genes that are differentially regulated between diploid and autotetraploid piNs in both humans and chimpanzees, suggesting that the tetraploid state leads to specific gene expression changes. This could be due to the lower plating density for tetraploid piNs (Shan et al., 2024), due to the larger size of their soma, or to differences in the relative soma to neurite ratio in tetraploid cells. Nevertheless, the genes that we focus on in this study are *cis*-regulated in allotetraploids where the effects of the cellular environment are controlled for, and our follow-up studies on these genes are all done in diploid piNs.

### ATAC-seq analysis

We aligned ATAC-seq data from human and chimpanzee diploid, autotetraploid, and allotetraploid piNs to a composite human-chimpanzee genome (from hg38 and panTro6) with bwa mem v0.7.17 (Li and Durbin, 2009). We filtered poorly aligning reads and duplicates using samtools (samtools view -F 3844 -q 10) (Li and Durbin, 2009). We made bed files from bam files using bedtools v2.26.0 (Quinlan and Hall, 2010). We used MACS2 v2.1.1.201160309 to call peaks (–nomodel –extsize 200) (Zhang et al., 2008). We used regions where reads from diploid and autotetraploid piNs mapped to the incorrect species as the background for peak calling. We removed data from one piN line for downstream analysis due to poor ATAC-seq data quality: H1H1a_H6-1.

We filtered peaks and reads by mapping reads from panTro6 to hg38 and vice versa with pslMap (Zhu et al., 2007). The chain file panTro6.hg38.all.chain.gz from UCSC (Navarro Gonzalez et al., 2021) was used in pslMap after filtering for orthologous chains as previously described (Turakhia et al., 2020). We filtered for uniquely mapping reads and peaks where orthologous regions have <5-fold change and <5kb size difference. We removed peaks that overlap ENCODE blacklist regions in hg38 (Amemiya et al., 2019) and alternate chromosomes. We generated reads and peaks listed in hg38 and panTro6 coordinates for each sample. Reads that mapped to hg38 initially in allotetraploids were represented separately than those that mapped to panTro6.

To generate a background mis-mapping file in hg38 and panTro6 coordinates for both species, we used reads from diploids and autotetraploids that mapped to the incorrect species. Using these mis-mapping files, we called peaks using MACS2 to use as a background set of genomic regions where there were high levels of mis-mapping.

We performed allele-specific accessibility (ASA) and differential accessibility (DA) analysis with DiffBind v3.0.15 (Ross-Innes et al., 2012) for reads and peaks in hg38 and panTro6 co-ordinates separately using the background mis-mapping files. To identify the ATAC peaks for analysis with Diffbind, we took the top 30,000 peaks per sample by macs2 q-value and identified peaks found in at least three samples and at least two different lines. This resulted in a set of 84,945 ATAC peaks. 57,330 of these ATAC peaks did not overlap coding regions. We called peaks as significant if they had an adjusted p<0.05 after Benjamini-Hochberg FDR correction. We kept peaks only if they had the same direction-of-effect when assessed in both hg38 and panTro6 coordinates.

We classified peaks by regulatory type based on the following: *cis*: significant DA, significant ASA, Log_2_(FC) in the same direction between ASA and DA. *trans*: significant DA, not significant ASA. *compensatory*: significant ASA, not significant DA.

Differential accessibility was also assessed between KCl timepoints by comparing stimulated to unstimulated piN samples. We excluded peaks that overlapped peaks called from background mis-mapping files in all downstream analyses. We also removed ATAC peaks that overlap chr21:43377000-43584000 where there is an assembly error in hg38.

Motif enrichment was calculated for ATAC-seq peaks that were significantly induced with KCl (30min or 2hr) and *cis*-regulated using HOMER (Heinz et al., 2010) findMotifsGenome.pl. *De novo* motifs are reported.

### CUT&Tag analysis

CUT&Tag data was processed as described for ATAC-seq with some minor differences. Before mapping, we used cutadapt v2.5 (Martin, 2011) to trim adapters. MACS2 was used to call peaks using matched IgG samples as the background. The following parameters were used for peak calling: –nomodel –shift -100 –extsize 200. Peaks assessed with DiffBind had to be found in at least 3 samples.

### Variant Analysis within FOS CUT&Tag peaks

Filtered net files for syntenic alignments in maf format (*synNet.maf) were downloaded from the UCSC Genome Browser between hg38 and panTro6, gorGor6, rheMac10, and calJac4 and parsed for variants between the human genome (hg38) and each primate species. Regions of the human genome without a syntenic alignment were considered missing data. For human-biased, chimpanzee-biased, and unbiased CUT&Tag peaks, bedtools intersect was used to intersect variant locations with peak locations. To identify variants within AP-1 and CUX2 motifs, HOMER (Heinz et al., 2010) annotatePeaks.pl was used with CUX2 and AP-1 motif annotation files. Motif locations from HOMER were then intersected with variant locations to find variants within motifs. Variant frequency within a motif was calculated as the number of variants overlapping the motif divided by the number of instances of that motif multiplied by the motif length.

## References

Abrahams, B. S., Arking, D. E., Campbell, D. B., Mefford, H. C., Morrow, E. M., Weiss, L. A., Menashe, I., Wadkins, T., Banerjee-Basu, S., and Packer, A., et al., 2013. Sfari gene 2.0: a community-driven knowledgebase for the autism spectrum disorders (asds). Molecular Autism, 4(1):36.

Agoglia, R. M., Sun, D., Birey, F., Yoon, S.-J., Miura, Y., Sabatini, K., Pasca, S. P., and Fraser, H. B., 2021. Primate cell fusion disentangles gene regulatory divergence in neurodevelopment. Nature, 592(7854):421–427.

Amemiya, H. M., Kundaje, A., and Boyle, A. P., 2019. The encode blacklist: Identification of problematic regions of the genome. Scientific Reports, 9(1):9354.

Ataman, B., Boulting, G. L., Harmin, D. A., Yang, M. G., Baker-Salisbury, M., Yap, E.-L., Malik, A. N., Mei, K., Rubin, A. A., Spiegel, I., et al., 2016. Evolution of osteocrin as an activity-regulated factor in the primate brain. Nature, 539(7628):242–247.

Atsaves, V., Leventaki, V., Rassidakis, G. Z., and Claret, F. X., 2019. Ap-1 transcription factors as regulators of immune responses in cancer. Cancers, 11(77):1037.

Bading, H., Ginty, D. D., and Greenberg, M. E., 1993. Regulation of gene expression in hippocampal neurons by distinct calcium signaling pathways. Science, 260(5105):181–186.

Baum, M. L., Wilton, D. K., Fox, R. G., Carey, A., Hsu, Y.-H. H., Hu, R., Jäntti, H. J., Fahey, J. B., Muthukumar, A. K., Salla, N., et al., 2024. Csmd1 regulates brain complement activity and circuit development. Brain, Behavior, and Immunity, 119:317–332.

Boulting, G. L., Durresi, E., Ataman, B., Sherman, M. A., Mei, K., Harmin, D. A., Carter, A. C., Hochbaum, D. R., Granger, A. J., Engreitz, J. M., et al., 2021. Activity-dependent regulome of human gabaergic neurons reveals new patterns of gene regulation and neurological disease heritability. Nature Neuroscience, 24(3):437–448.

Buenrostro, J. D., Giresi, P. G., Zaba, L. C., Chang, H. Y., and Greenleaf, W. J., 2013. Transposition of native chromatin for fast and sensitive epigenomic profiling of open chromatin, dna-binding proteins and nucleosome position. Nature Methods, 10(12):1213–1218.

Byrns, C. N., Saikumar, J., and Bonini, N. M., 2021. Glial ap1 is activated with aging and accelerated by traumatic brain injury. Nature Aging, 1(7):585–597.

Caglayan, E., Ayhan, F., Liu, Y., Vollmer, R. M., Oh, E., Sherwood, C. C., Preuss, T. M., Yi, S. V., and Konopka, G., 2023. Molecular features driving cellular complexity of human brain evolution. Nature, 620(7972):145–153.

Carey, M. B. and Matsumoto, S. G., 1999. Spontaneous calcium transients are required for neuronal differentiation of murine neural crest. Developmental Biology, 215(2):298–313.

Carroll, S. B., 2008. Evo-devo and an expanding evolutionary synthesis: A genetic theory of morphological evolution. Cell, 134(1):25–36.

Ciceri, G., Baggiolini, A., Cho, H. S., Kshirsagar, M., Benito-Kwiecinski, S., Walsh, R. M., Aromolaran, K. A., Gonzalez-Hernandez, A.J., Munguba, H., Koo, S. Y., et al., 2024. An epigenetic barrier sets the timing of human neuronal maturation. Nature, 626(8000):881–890.

Corces, M. R., Trevino, A. E., Hamilton, E. G., Greenside, P. G., Sinnott-Armstrong, N. A., Vesuna, S., Satpathy, A. T., Rubin, A. J., Montine, K. S., Wu, B., et al., 2017. An improved atac-seq protocol reduces background and enables interrogation of frozen tissues. Nature Methods, 14(10):959–962.

Countryman, R. A., Kaban, N. L., and Colombo, P. J., 2005. Hippocampal c-fos is necessary for long-term memory of a socially transmitted food preference. Neurobiology of Learning and Memory, 84(3):175–183.

Courchet, J., Lewis, T. L., Lee, S., Courchet, V., Liou, D.-Y., Aizawa, S., and Polleux, F., 2013. Terminal axon branching is regulated by the lkb1-nuak1 kinase pathway via presynaptic mitochondrial capture. Cell, 153(7):1510–1525.

Creyghton, M. P., Cheng, A. W., Welstead, G. G., Kooistra, T., Carey, B. W., Steine, E. J., Hanna, J., Lodato, M. A., Frampton, G. M., Sharp, P. A., et al., 2010. Histone h3k27ac separates active from poised enhancers and predicts developmental state. Proceedings of the National Academy of Sciences of the United States of America, 107(50):21931–21936.

Cubelos, B., Sebastián-Serrano, A., Beccari, L., Calcagnotto, M. E., Cisneros, E., Kim, S., Dopazo, A., Alvarez-Dolado, M., Redondo, J. M., Bovolenta, P., et al., 2010. Cux1 and cux2 regulate dendritic branching, spine morphology, and synapses of the upper layer neurons of the cortex. Neuron, 66(4):523–535.

Dardou, D., Monlezun, S., Foerch, P., Courade, J. P., Cuvelier, L., De Ryck, M., and Schiffmann, S. N., 2013. A role for sv2c in basal ganglia functions. Brain Research, 1507:61–73.

Deisseroth, K., Singla, S., Toda, H., Monje, M., Palmer, T. D., and Malenka, R. C., 2004. Excitation-neurogenesis coupling in adult neural stem/progenitor cells. Neuron, 42(4):535–552.

Dobin, A., Davis, C. A., Schlesinger, F., Drenkow, J., Zaleski, C., Jha, S., Batut, P., Chaisson, M., and Gingeras, T. R., 2013. Star: ultrafast universal rna-seq aligner. Bioinformatics (Oxford, England), 29(1):15–21.

Doench, J. G., Fusi, N., Sullender, M., Hegde, M., Vaimberg, E. W., Donovan, K. F., Smith, I., Tothova, Z., Wilen, C., Orchard, R., et al., 2016. Optimized sgrna design to maximize activity and minimize off-target effects of crispr-cas9. Nature Biotechnology, 34(2):184–191.

Dunn, A. R., Stout, K. A., Ozawa, M., Lohr, K. M., Hoffman, C. A., Bernstein, A. I., Li, Y., Wang, M., Sgobio, C., Sastry, N., et al., 2017. Synaptic vesicle glycoprotein 2c (sv2c) modulates dopamine release and is disrupted in parkinson disease. Proceedings of the National Academy of Sciences, 114(11):E2253–E2262.

Espuny-Camacho, I., Michelsen, K. A., Gall, D., Linaro, D., Hasche, A., Bonnefont, J., Bali, C., Orduz, D., Bilheu, A., Herpoel, A., et al., 2013. Pyramidal neurons derived from human pluripotent stem cells integrate efficiently into mouse brain circuits in vivo. Neuron, 77(3):440–456.

Fair, S. R., Julian, D., Hartlaub, A. M., Pusuluri, S. T., Malik, G., Summerfied, T. L., Zhao, G., Hester, A. B., Ackerman, W. E., Hollingsworth, E. W., et al., 2020. Electrophysiological maturation of cerebral organoids correlates with dynamic morphological and cellular development. Stem Cell Reports, 15(4):855–868.

Faria-Pereira, A., Temido-Ferreira, M., and Morais, V. A., 2022. Brainphys neuronal media support physiological function of mitochondria in mouse primary neuronal cultures. Frontiers in Molecular Neuroscience, 15.

Favuzzi, E., Marques-Smith, A., Deogracias, R., Winterflood, C. M., Sánchez-Aguilera, A., Mantoan, L., Maeso, P., Fernandes, C., Ewers, H., and Rico, B., et al., 2017. Activity-dependent gating of parvalbumin interneuron function by the perineuronal net protein brevican. Neuron, 95(3):639– 655.e10.

Fischer-Colbrie, R., Laslop, A., and Kirchmair, R., 1995. Secretogranin ii: molecular properties, regulation of biosynthesis and processing to the neuropeptide secretoneurin. Progress in Neurobiology, 46(1):49–70.

Gallego Romero, I., Pavlovic, B. J., Hernando-Herraez, I., Zhou, X., Ward, M. C., Banovich, N. E., Kagan, C. L., Burnett, J. E., Huang, C. H., Mitrano, A., et al., 2015. A panel of induced pluripotent stem cells from chimpanzees: a resource for comparative functional genomics. eLife, 4:e07103.

Gaspard, N., Bouschet, T., Hourez, R., Dimidschstein, J., Naeije, G., van den Ameele, J., Espuny-Camacho, I., Herpoel, A., Passante, L., Schiffmann, S. N., et al., 2008. An intrinsic mechanism of corticogenesis from embryonic stem cells. Nature, 455(7211):351–357.

Glauser, D. A. and Schlegel, W., 2007. Sequential actions of erk1/2 on the ap-1 transcription factor allow temporal integration of metabolic signals in pancreatic beta cells. FASEB journal: official publication of the Federation of American Societies for Experimental Biology, 21(12):3240–3249.

Gokhman, D., Agoglia, R. M., Kinnebrew, M., Gordon, W., Sun, D., Bajpai, V. K., Naqvi, S., Chen, C., Chan, A., Chen, C., et al., 2021. Human-chimpanzee fused cells reveal cisregulatory divergence underlying skeletal evolution. Nature Genetics, 53(4):467–476.

Greenberg, M. E. and Ziff, E. B., 1984. Stimulation of 3t3 cells induces transcription of the c-fos proto-oncogene. Nature, 311(5985):433–438.

Gu, Y., Janoschka, S., and Ge, S., 2012. Neurogenesis and Hippocampal Plasticity in Adult Brain, volume 15 of Current Topics in Behavioral Neurosciences, page 31–48. Springer Berlin Heidelberg, Berlin, Heidelberg.

He, J., Yamada, K., and Nabeshima, T., 2002. A role of fos expression in the ca3 region of the hippocampus in spatial memory formation in rats. Neuropsychopharmacology, 26(2):259–268.

Heinz, S., Benner, C., Spann, N., Bertolino, E., Lin, Y. C., Laslo, P., Cheng, J. X., Murre, C., Singh, H., and Glass, C. K., et al., 2010. Simple combinations of lineage-determining transcription factors prime cis-regulatory elements required for macrophage and b cell identities. Molecular Cell, 38(4):576–589.

Hergenreder, E., Minotti, A. P., Zorina, Y., Oberst, P., Zhao, Z., Munguba, H., Calder, E. L., Baggiolini, A., Walsh, R. M., Liston, C., et al., 2024. Combined small-molecule treatment accelerates maturation of human pluripotent stem cell-derived neurons. Nature Biotechnology, 42(10):1515–1525.

Herring, C. A., Simmons, R. K., Freytag, S., Poppe, D., Moffet, J. J. D., Pflueger, J., Buckberry, S., Vargas-Landin, D. B., Clément, O., Echeverría, E. G., et al., 2022. Human prefrontal cortex gene regulatory dynamics from gestation to adulthood at single-cell resolution. Cell, 185(23):4428– 4447.e28.

Holliday, J., Adams, R. J., Sejnowski, T. J., and Spitzer, N. C., 1991. Calcium-induced release of calcium regulates differentiation of cultured spinal neurons. Neuron, 7(5):787–796.

Horlbeck, M. A., Gilbert, L. A., Villalta, J. E., Adamson, B., Pak, R. A., Chen, Y., Fields, A. P., Park, C. Y., Corn, J. E., Kampmann, M., et al., 2016. Compact and highly active next-generation libraries for crispr-mediated gene repression and activation. eLife, 5:e19760.

Hrvatin, S., Hochbaum, D. R., Nagy, M. A., Cicconet, M., Robertson, K., Cheadle, L., Zilionis, R., Ratner, A., Borges-Monroy, R., Klein, A. M., et al., 2018. Single-cell analysis of experience-dependent transcriptomic states in the mouse visual cortex. Nature Neuroscience, 21(1):120–129.

Iqbal, Z., Willemsen, M. H., Papon, M.-A., Musante, L., Benevento, M., Hu, H., Venselaar, H., Wissink-Lindhout, W. M., Vulto-van Silfhout, A. T., Vissers, L. E. L. M., et al., 2015. Homozygous slc6a17 mutations cause autosomal-recessive intellectual disability with progressive tremor, speech impairment, and behavioral problems. The American Journal of Human Genetics, 96(3):386–396.

Iwata, R., Casimir, P., Erkol, E., Boubakar, L., Planque, M., Gallego López, I. M., Ditkowska, M., Gaspariunaite, V., Beckers, S., Remans, D., et al., 2023. Mitochondria metabolism sets the species-specific tempo of neuronal development. Science, 379(6632):eabn4705.

Jorstad, N. L., Song, J. H. T., Exposito-Alonso, D., Suresh, H., Castro-Pacheco, N., Krienen, F. M., Yanny, A. M., Close, J., Gelfand, E., Long, B., et al., 2023. Comparative transcriptomics reveals human-specific cortical features. Science (New York, N.Y.), 382(6667):eade9516.

Kaya-Okur, H. S., Wu, S. J., Codomo, C. A., Pledger, E. S., Bryson, T. D., Henikoff, J. G., Ahmad, K., and Henikoff, S., 2019. Cut&tag for efficient epigenomic profiling of small samples and single cells. Nature Communications, 10(1):1930.

King, M.-C. and Wilson, A. C., 1975. Evolution at two levels in humans and chimpanzees: Their macromolecules are so alike that regulatory mutations may account for their biological differences. Science, 188(4184):107–116.

Koopmans, F., van Nierop, P., Andres-Alonso, M., Byrnes, A., Cijsouw, T., Coba, M. P., Cornelisse, L. N., Farrell, R. J., Goldschmidt, H. L., Howrigan, D. P., et al., 2019. Syngo: An evidence-based, expert-curated knowledge base for the synapse. Neuron, 103(2):217–234.e4.

Lanfranchi, M., Yandiev, S., Meyer-Dilhet, G., Ellouze, S., Kerkhofs, M., Dos Reis, R., Garcia, A., Blondet, C., Amar, A., Kneppers, A., et al., 2024. The ampk-related kinase nuak1 controls cortical axons branching by locally modulating mitochondrial metabolic functions. Nature Communications, 15(1):2487.

Leblond, C. S., Le, T.-L., Malesys, S., Cliquet, F., Tabet, A.-C., Delorme, R., Rolland, T., and Bourgeron, T., 2021. Operative list of genes associated with autism and neurodevelopmental disorders based on database review. Molecular and Cellular Neuroscience, 113:103623.

Li, H. and Durbin, R., 2009. Fast and accurate short read alignment with burrows–wheeler transform. Bioinformatics, 25(14):1754–1760.

Liao, Y., Smyth, G. K., and Shi, W., 2014. featurecounts: an efficient general purpose program for assigning sequence reads to genomic features. Bioinformatics, 30(7):923–930.

Lin, Y., Bloodgood, B. L., Hauser, J. L., Lapan, A. D., Koon, A. C., Kim, T.-K., Hu, L. S., Malik, A. N., and Greenberg, M. E., 2008. Activity-dependent regulation of inhibitory synapse development by npas4. Nature, 455(7217):1198–1204.

Lindhout, F. W., Szafranska, H. M., Imaz-Rosshandler, I., Guglielmi, L., Moarefian, M., Voitiuk, K., Zernicka-Glover, N. K., Boulanger, J., Schulze, U., Sánchez, D. J. L.D., et al., 2025. Calcium dynamics tune developmental tempo to generate evolutionarily divergent axon tract lengths.:2024.12.28.630576.

Liu, J., Li, S., Li, X., Li, W., Yang, Y., Guo, S., Lv, L., Xiao, X., Yao, Y.-G., Guan, F., et al., 2021. Genome-wide association study followed by trans-ancestry meta-analysis identify 17 new risk loci for schizophrenia. BMC medicine, 19(1):177.

Love, M. I., Huber, W., and Anders, S., 2014. Moderated estimation of fold change and dispersion for rna-seq data with deseq2. Genome Biology, 15(12):550.

Malik, A. N., Vierbuchen, T., Hemberg, M., Rubin, A. A., Ling, E., Couch, C. H., Stroud, H., Spiegel, I., Farh, K. K.-H., Harmin, D. A., et al., 2014. Genome-wide identification and characterization of functional neuronal activity-dependent enhancers. Nature Neuroscience, 17(10):1330–1339.

Martin, M., 2011. Cutadapt removes adapter sequences from high-throughput sequencing reads. EMBnet.journal, 17(11):10–12.

Martínez-Zamudio, R. I., Roux, P.-F., de Freitas, J. A. N. L. F., Robinson, L., Doré, G., Sun, B., Belenki, D., Milanovic, M., Herbig, U., Schmitt, C. A., et al., 2020. Ap-1 imprints a reversible transcriptional programme of senescent cells. Nature Cell Biology, 22(7):842–855.

McLean, C. Y., Reno, P. L., Pollen, A. A., Bassan, A. I., Capellini, T. D., Guenther, C., Indjeian, V. B., Lim, X., Menke, D. B., Schaar, B. T., et al., 2011. Human-specific loss of regulatory dna and the evolution of human-specific traits. Nature, 471(7337):216–219.

Napoli, A. and Obeid, I., 2015. Investigating brain functional evolution and plasticity using microelectrode array technology. Brain Research Bulletin, 119:127–135.

Navarro Gonzalez, J., Zweig, A. S., Speir, M. L., Schmelter, D., Rosenbloom, K. R., Raney, B. J., Powell, C. C., Nassar, L. R., Maulding, N. D., Lee, C. M., et al., 2021. The ucsc genome browser database: 2021 update. Nucleic Acids Research, 49(D1):D1046–D1057.

Nehme, R., Zuccaro, E., Ghosh, S. D., Li, C., Sherwood, J. L., Pietilainen, O., Barrett, L. E., Limone, F., Worringer, K. A., Kommineni, S., et al., 2018. Combining ngn2 programming with developmental patterning generates human excitatory neurons with nmdar-mediated synaptic transmission. Cell Reports, 23(8):2509–2523.

Nieto, M., Monuki, E. S., Tang, H., Imitola, J., Haubst, N., Khoury, S. J., Cunningham, J., Gotz, M., and Walsh, C. A., 2004. Expression of cux-1 and cux-2 in the subventricular zone and upper layers ii-iv of the cerebral cortex. The Journal of Comparative Neurology, 479(2):168–180.

Ordureau, A., Paulo, J. A., Zhang, J., An, H., Swatek, K. N., Cannon, J. R., Wan, Q., Komander, D., and Harper, J. W., 2020. Global landscape and dynamics of parkin and usp30dependent ubiquitylomes in ineurons during mitophagic signaling. Molecular Cell, 77(5):1124–1142.e10.

Otani, T., Marchetto, M. C., Gage, F. H., Simons, B. D., and Livesey, F. J., 2016. 2d and 3d stem cell models of primate cortical development identify species-specific differences in progenitor behavior contributing to brain size. Cell Stem Cell, 18(4):467–480.

Patrick, R., Naval-Sanchez, M., Deshpande, N., Huang, Y., Zhang, J., Chen, X., Yang, Y., Tiwari, K., Esmaeili, M., Tran, M., et al., 2024. The activity of early-life gene regulatory elements is hijacked in aging through pervasive ap-1-linked chromatin opening. Cell Metabolism, 36(8):1858–1881.e23.

Paylor, R., Johnson, R. S., Papaioannou, V., Spiegelman, B. M., and Wehner, J. M., 1994. Behavioral assessment of c-fos mutant mice. Brain Research, 651(1):275–282.

Piatti, V. C., Davies-Sala, M. G., Espósito, M. S., Mongiat, L. A., Trinchero, M. F., and Schinder, A. F., 2011. The timing for neuronal maturation in the adult hippocampus is modulated by local network activity. The Journal of Neuroscience: The Official Journal of the Society for Neuroscience, 31(21):7715–7728.

Pollina, E. A., Gilliam, D. T., Landau, A. T., Lin, C., Pajarillo, N., Davis, C. P., Harmin, D. A., Yap, E.-L., Vogel, I. R., Griffith, E. C., et al., 2023. A npas4–nua4 complex couples synaptic activity to dna repair. Nature, 614(7949):732–741.

Pruunsild, P., Bengtson, C. P., and Bading, H., 2017. Networks of cultured ipsc-derived neurons reveal the human synaptic activity-regulated adaptive gene program. Cell Reports, 18(1):122–135.

Qiu, J., McQueen, J., Bilican, B., Dando, O., Magnani, D., Punovuori, K., Selvaraj, B. T., Livesey, M., Haghi, G., Heron, S., et al., 2016. Evidence for evolutionary divergence of activity-dependent gene expression in developing neurons. eLife, 5:e20337.

Quinlan, A. R. and Hall, I. M., 2010. Bedtools: a flexible suite of utilities for comparing genomic features. Bioinformatics, 26(6):841–842.

Replogle, J. M., Norman, T. M., Xu, A., Hussmann, J. A., Chen, J., Cogan, J. Z., Meer, E. J., Terry, J. M., Riordan, D. P., Srinivas, N., et al., 2020. Combinatorial single-cell crispr screens by direct guide rna capture and targeted sequencing. Nature Biotechnology, 38(8):954–961.

Ringler, S. L., Aye, J., Byrne, E., Anderson, M., and Turner, C. P., 2008. Effects of disrupting calcium homeostasis on neuronal maturation: Early inhibition and later recovery. Cellular and Molecular Neurobiology, 28(3):389–409.

Ross-Innes, C. S., Stark, R., Teschendorff, A. E., Holmes, K. A., Ali, H. R., Dunning, M. J., Brown, G. D., Gojis, O., Ellis, I. O., Green, A. R., et al., 2012. Differential oestrogen receptor binding is associated with clinical outcome in breast cancer. Nature, 481(7381):389–393.

Sanchez-Priego, C., Hu, R., Boshans, L. L., Lalli, M., Janas, J. A., Williams, S. E., Dong, Z., and Yang, N., 2022. Mapping cis-regulatory elements in human neurons links psychiatric disease heritability and activity-regulated transcriptional programs. Cell Reports, 39(9).

Sanson, K. R., Hanna, R. E., Hegde, M., Donovan, K. F., Strand, C., Sullender, M. E., Vaimberg, E. W., Goodale, A., Root, D. E., Piccioni, F., et al., 2018. Optimized libraries for crispr-cas9 genetic screens with multiple modalities. Nature Communications, 9(1):5416.

Schivell, A. E., Mochida, S., Kensel-Hammes, P., Custer, K. L., and Bajjalieh, S. M., 2005. Sv2a and sv2c contain a unique synaptotagmin-binding site. Molecular and Cellular Neuroscience, 29(1):56–64.

Senís, E., Esgleas, M., Najas, S., Jiménez-Sábado, V., Bertani, C., Giménez-Alejandre, M., Escriche, A., Ruiz-Orera, J., Hergueta-Redondo, M., Jiménez, M., et al., 2021. Tunar lncrna encodes a microprotein that regulates neural differentiation and neurite formation by modulating calcium dynamics. Frontiers in Cell and Developmental Biology, 9.

Shan, X., Zhang, A., Rezzonico, M. G., Tsai, M.-C., Sanchez-Priego, C., Zhang, Y., Chen, M. B., Choi, M., Andrade López, J. M., Phu, L., et al., 2024. Fully defined ngn2 neuron protocol reveals diverse signatures of neuronal maturation. Cell Reports Methods, 4(9):100858.

Sheng, M., Dougan, S. T., McFadden, G.,, and Greenberg, M. E., 1988. Calcium and growth factor pathways of c-fos transcriptional activation require distinct upstream regulatory sequences. Molecular and Cellular Biology, 8(7):2787–2796.

Sheng, M. and Greenberg, M. E., 1990. The regulation and function of c-fos and other immediate early genes in the nervous system. Neuron, 4(4):477–485.

Song, J. H., Carter, A. C., Bushinsky, E. M., Beck, S. G., Petrocelli, J. E., Koreman, G. T., Babu, J., Kingsley, D. M., Greenberg, M. E., and Walsh, C. A., et al., 2025. Human-chimpanzee tetraploid system defines mechanisms of species-specific neural gene regulation. Submitted,.

Song, J. H. T., Grant, R. L., Behrens, V. C., Kučka, M., Roberts Kingman, G.A., Soltys, V., Chan, Y. F., and Kingsley, D. M., 2021. Genetic studies of human-chimpanzee divergence using stem cell fusions. Proceedings of the National Academy of Sciences of the United States of America, 118(51):e2117557118.

Stroud, H., Yang, M. G., Tsitohay, Y. N., Davis, C. P., Sherman, M. A., Hrvatin, S., Ling, E., and Greenberg, M. E., 2020. An activity-mediated transition in transcription in early postnatal neurons. Neuron, 107(5):874–890.e8.

Tao, X., Finkbeiner, S., Arnold, D. B., Shaywitz, A. J., and Greenberg, M. E., 1998. Ca2+ influx regulates bdnf transcription by a creb family transcription factor-dependent mechanism. Neuron, 20(4):709–726.

Tian, R., Gachechiladze, M. A., Ludwig, C. H., Laurie, M. T., Hong, J. Y., Nathaniel, D., Prabhu, A. V., Fernandopulle, M. S., Patel, R., Abshari, M., et al., 2019. Crispr interference-based platform for multimodal genetic screens in human ipsc-derived neurons. Neuron, 104(2):239– 255.e12.

Traunmüller, L., Duffy, E. E., Liu, H., Sanalidou, S., Assad, E. G., Sun, S., Pajarillo, N. S., Niu, N., Griffith, E. C., and Greenberg, M. E., et al., 2025. Novel environment exposure drives temporally defined and region-specific chromatin accessibility and gene expression changes in the hippocampus.:2024.10.31.621351.

Turakhia, Y., Maio, N. D., Thornlow, B., Gozashti, L., Lanfear, R., Walker, C. R., Hinrichs, A. S., Fernandes, J. D., Borges, R., Slodkowicz, G., et al., 2020. Stability of sars-cov-2 phylogenies. PLOS Genetics, 16(11):e1009175.

Vanderhaeghen, P. and Polleux, F., 2023. Developmental mechanisms underlying the evolution of human cortical circuits. Nature Reviews Neuroscience, 24(4):213–232.

Vierbuchen, T., Ling, E., Cowley, C. J., Couch, C. H., Wang, X., Harmin, D. A., Roberts, C. W. M., and Greenberg, M. E., 2017. Ap-1 transcription factors and the baf complex mediate signal-dependent enhancer selection. Molecular Cell, 68(6):1067–1082.e12.

Wagenaar, D. A., Pine, J., and Potter, S. M., 2006. Searching for plasticity in dissociated cortical cultures on multi-electrode arrays. Journal of Negative Results in BioMedicine, 5(1):16.

Wagner, E. F. and Eferl, R., 2005. Fos/ap-1 proteins in bone and the immune system. Immunological Reviews, 208(1):126–140.

Xia, Z., Dudek, H., Miranti, C. K., and Greenberg, M. E., 1996. Calcium influx via the nmda receptor induces immediate early gene transcription by a map kinase/erk-dependent mechanism. Journal of Neuroscience, 16(17):5425–5436.

Yang, M. G., Ling, E., Cowley, C. J., Greenberg, M. E., and Vierbuchen, T., 2022. Characterization of sequence determinants of enhancer function using natural genetic variation. eLife, 11:e76500.

Yap, E.-L. and Greenberg, M. E., 2018. Activity-regulated transcription: Bridging the gap between neural activity and behavior. Neuron, 100(2):330–348.

Yap, E.-L., Pettit, N. L., Davis, C. P., Nagy, M. A., Harmin, D. A., Golden, E., Dagliyan, O., Lin, C., Rudolph, S., Sharma, N., et al., 2021. Bidirectional perisomatic inhibitory plasticity of a fos neuronal network. Nature, 590(7844):115–121.

Zhang, Y., Liu, T., Meyer, C. A., Eeckhoute, J., Johnson, D. S., Bernstein, B. E., Nusbaum, C., Myers, R. M., Brown, M., Li, W., et al., 2008. Model-based analysis of chip-seq (macs). Genome Biology, 9(9):R137.

Zhu, J., Sanborn, J. Z., Diekhans, M., Lowe, C. B., Pringle, T. H., and Haussler, D., 2007. Comparative genomics search for losses of long-established genes on the human lineage. PLOS Computational Biology, 3(12):e247.

