## Supplemental Figures for "FOS binding sites are a hub for the evolution of activity-dependent gene regulatory programs in human neurons"

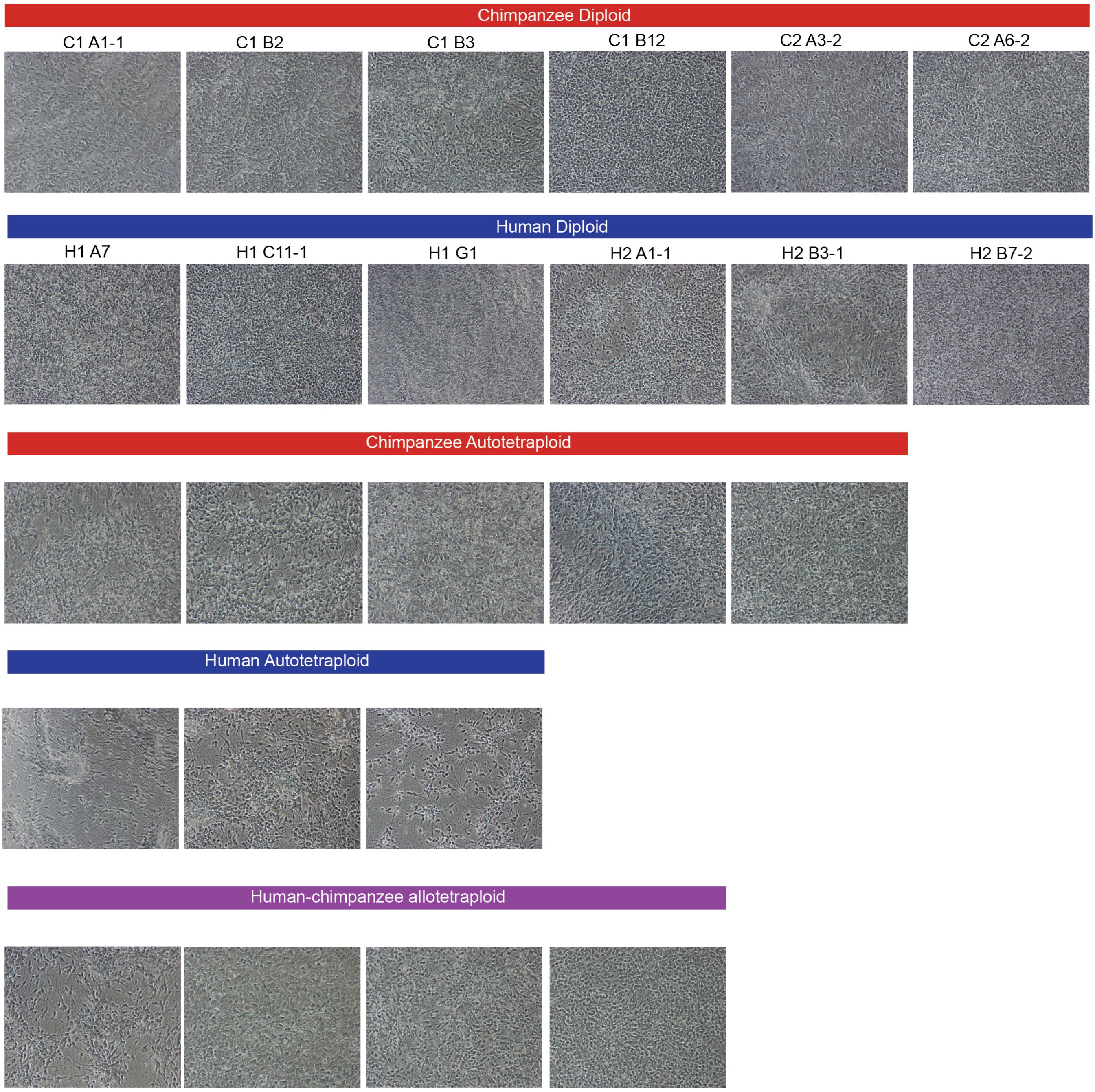


Figure S1: **Related to Figure 1** a. Brightfield images of piNs taken after 14 days of differentiation.

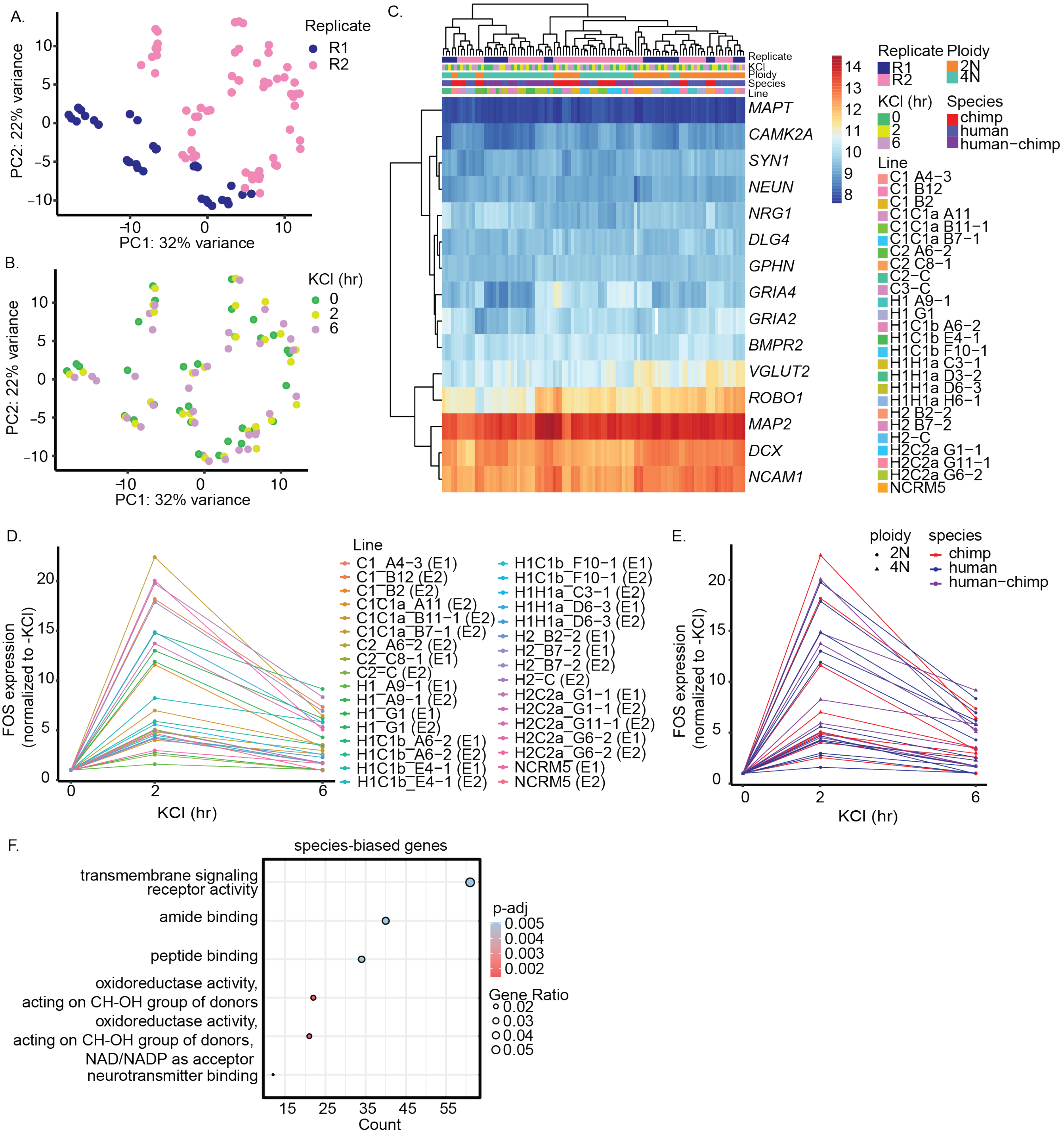


Figure S2: **Related to Figure 1** a. PCA plot for RNA-seq data showing PC1 and PC2 with samples colored by biological replicate. b. PCA plot for RNA-seq data showing PC1 and PC2 with samples colored by KCl stimulation condition. 0,2,6 refer to -KCl, 2hr KCl, and 6hr KCl treatment. c. Heatmap showing the expression levels of piN marker genes from (Nehme et al., 2018) in diploid and tetraploid piNs. d. Lineplot showing the expression of the IEG FOS at -KCl (0), 2hr KCl (2), and 6hr (6) time points for every cell line used in our initial RNA-seq analysis. Expression is normalized to the -KCl time point. e. Same as in d, but with the samples colored by species. 2N samples are indicated by circles, and 4N samples are indicated by triangles. f. GO analysis for species-biased genes (FC>1.5) in unstimulated piNs.

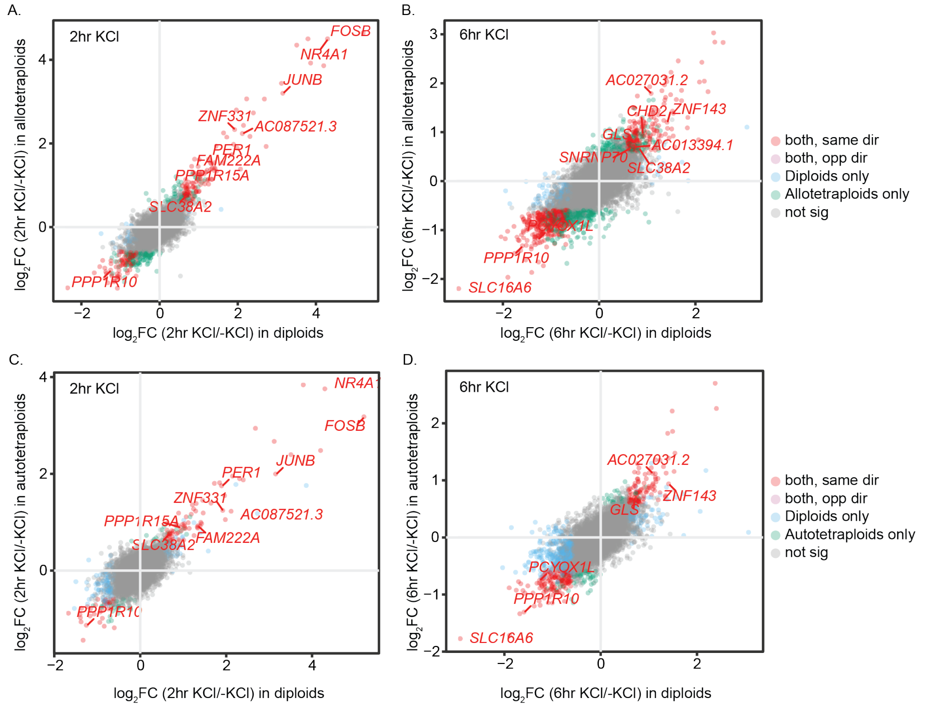


Figure S3: **Related to Figure 2.** a. Scatterplot showing the $\log_2$ fold change in expression between 2hr KCl and -KCl samples from allotetraploids vs the $\log_2$ fold change in expression between 2hr KCl and -KCl samples from diploids. Red points indicate genes that are activity-regulated in allotetraploids and diploids. Blue points indicate genes that are activity-regulated only in diploids. Green points indicate genes that are activity-regulated only in allotetraploids. b. Same as in a for 6hr KCl vs. -KCl samples. c. Same as in a for autotetraploids vs. diploids. d. Same as in b for autotetraploids vs. diploids.

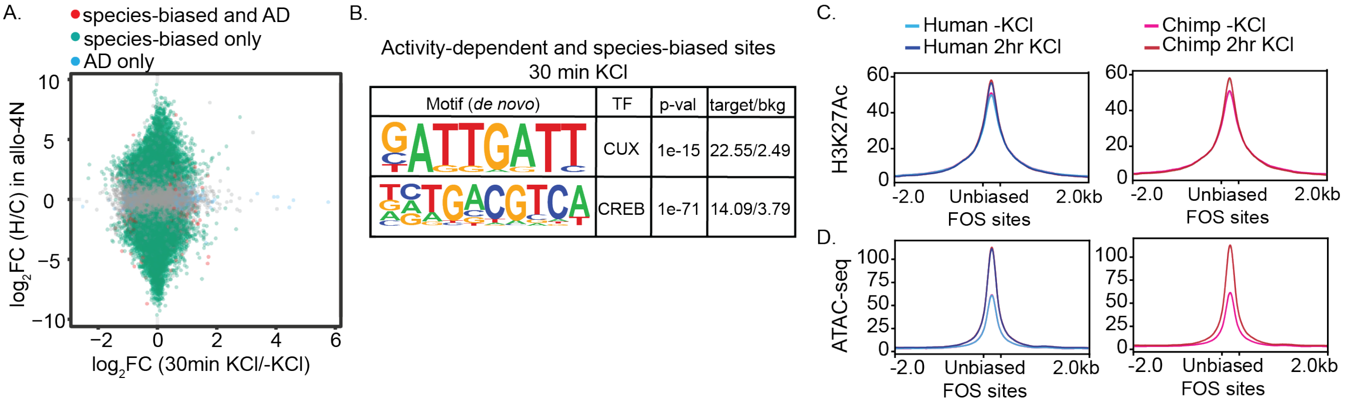


Figure S4: **Related to Figure 3.** a. Scatterplot showing the $\log_2$ fold change in allelic accessibility (human/chimpanzee) vs. the $\log_2$ fold change in accessibility with KCl treatment (30min KCl/-KCl) from ATAC-seq data. Red points indicate species-biased (SB) and activity-dependent (AD) sites, blue points indicate sites that are activity-dependent only, green points indicate sites that are species-biased only, and gray points are non-significant. b. *De novo* motif enrichment analysis at activity-dependent and species-biased ATAC-seq peaks after 30min KCl treatment. c. Average diagrams of H3K27Ac CUT&Tag signal at unbiased activity-dependent FOS binding sites. CUT&Tag data is from allotetraploids and signals from human and chimpanzee alleles are separated into two plots due to overlapping lines. d. Same as in c for ATAC-seq signal.

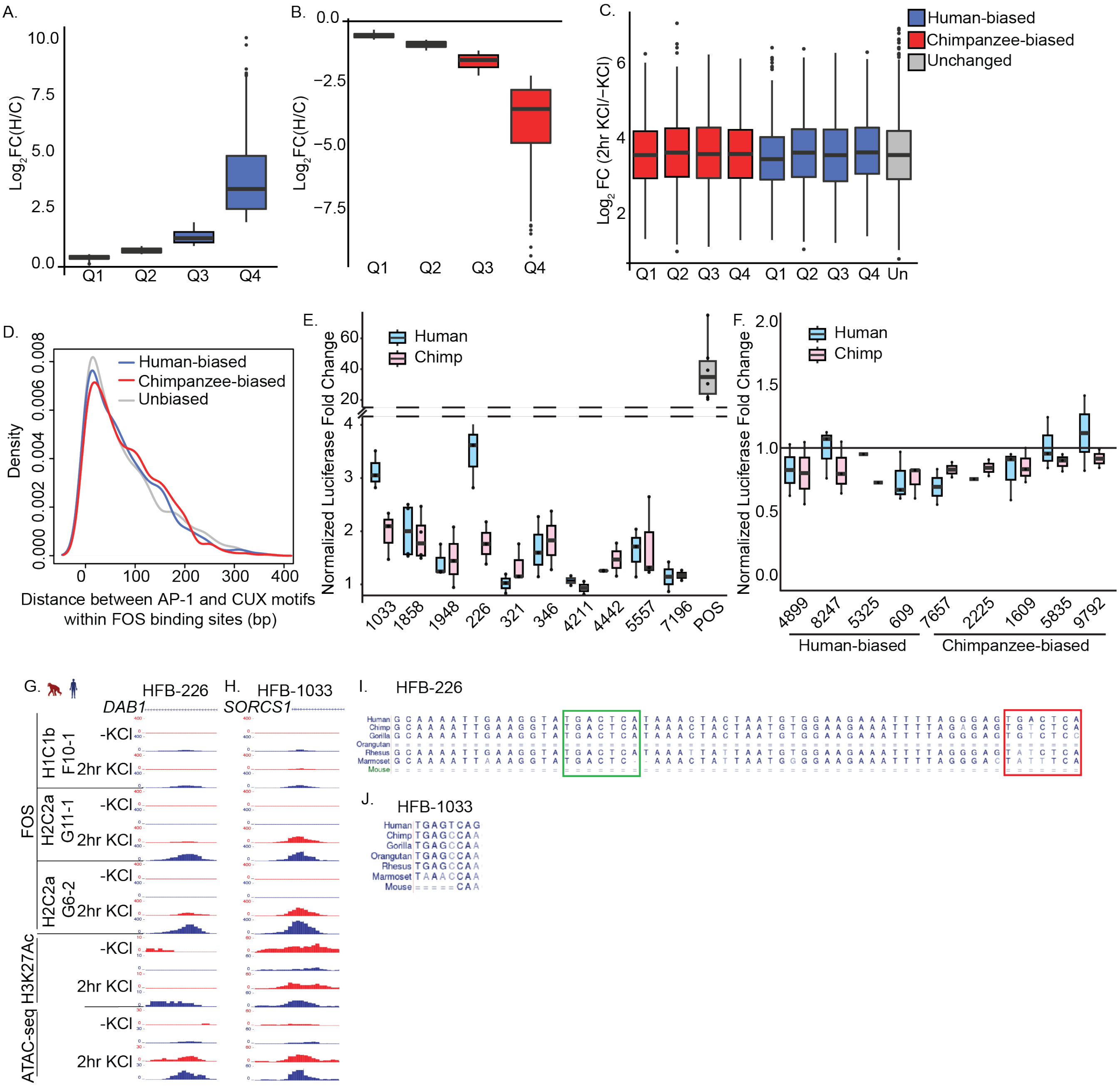


Figure S5: **Related to Figure 4.** a. Boxplot showing the $\log_2$ fold change in FOS binding between human and chimpanzee alleles for human-biased sites in quartiles 1-4. b. Same as in a for chimpanzee-biased sites. c. Boxplot showing the $\log_2$ fold change in FOS binding between 2hr KCl and minus KCl timepoints for human-biased and chimpanzee-biased quartiles 1-4 as well as all unbiased sites. d. Density plot showing the distribution of distances between AP-1 motifs and CUX motifs within FOS binding sites that are human-biased, chimpanzee-biased, or unbiased. The distance is only shown for peaks where AP-1 and CUX motifs occur within the same peak and the AP-1 sequence is unchanged between species. e. Boxplot of the normalized luciferase fold change (normalized to the empty negative control plasmid and to -KCl wells) for human-biased FOSbinding sites as well as the positive control from (Malik et al., 2014). f. Boxplot of the luciferase fold change for a second set of human-biased and chimpanzee-biased FOS binding sites tested in the luciferase assay. g. Tracks showing sequencing reads from the human and chimpanzee alleles at HBF-226. Displayed are tracks from three human-chimpanzee allotetraploid cell lines for FOS CUT&Tag at -KCl and 2hr KCl timepoints as well as H3K27Ac CUT&Tag and ATAC-seq for one human-chimpanzee allotetraploid cell line. Chimpanzee reads are depicted in red and human reads are depicted in blue. h. Same as in f for HBF-1033. i. Diagram showing the sequence of HBF-226 around two AP-1 motifs in human, chimpanzee, gorilla, orangutan, rhesus, marmoset, and mouse. The AP-1 motif boxed in green is intact in human and chimpanzee and was not changed in the luciferase assay. The AP-1 motif boxed in red was changed in the luciferase assay. j. Same as in i for the single AP-1 motif in HBF-1033.

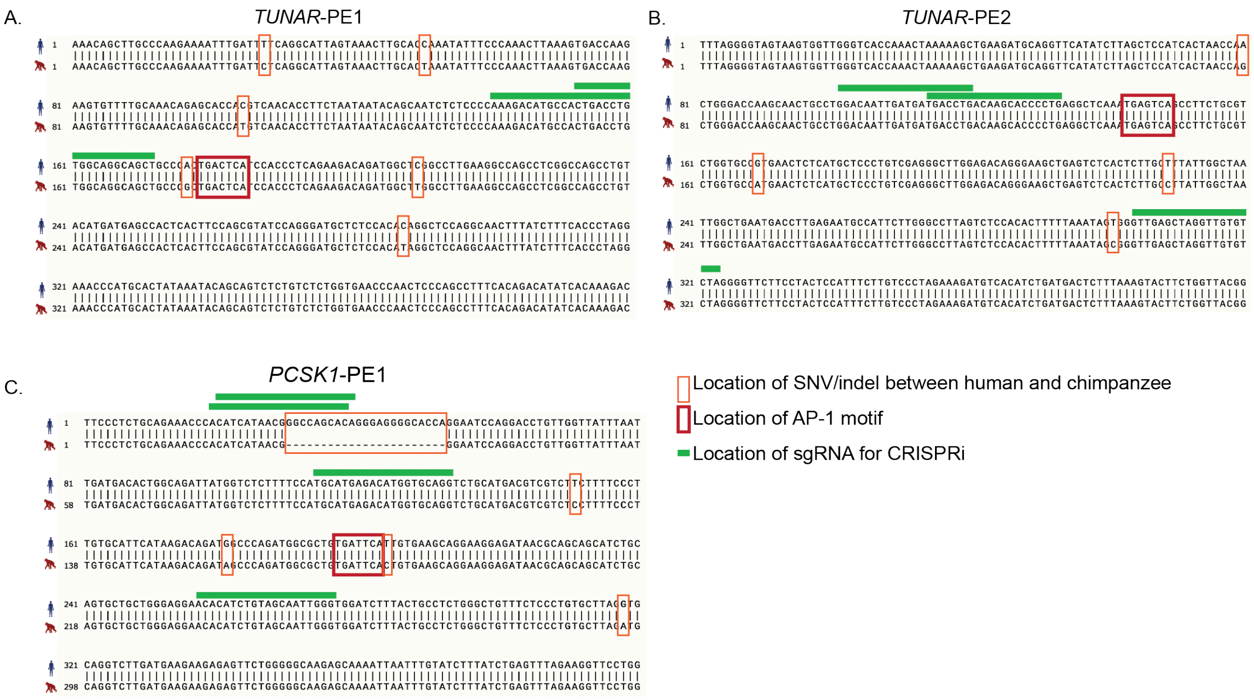


Figure S6: **Related to Figure 5.** a. Sequence alignment for *TUNAR*'s putative enhancer 1 (*TUNAR*-PE1) between human and chimpanzee. Human sequence is displayed on top and chimpanzee is displayed on bottom. The locations of SNVs/indels between the two species are indicated in orange boxes, AP-1 motifs are indicated in red boxes, and sgRNA sequences used for CRISPRi experiments are indicated as green bars. b. Same as in a for *TUNAR*'s putative enhancer 2 (*TUNAR*-PE2). c. Same as in a for *PCSK1*'s putative enhancer (*PCSK1*-PE1).

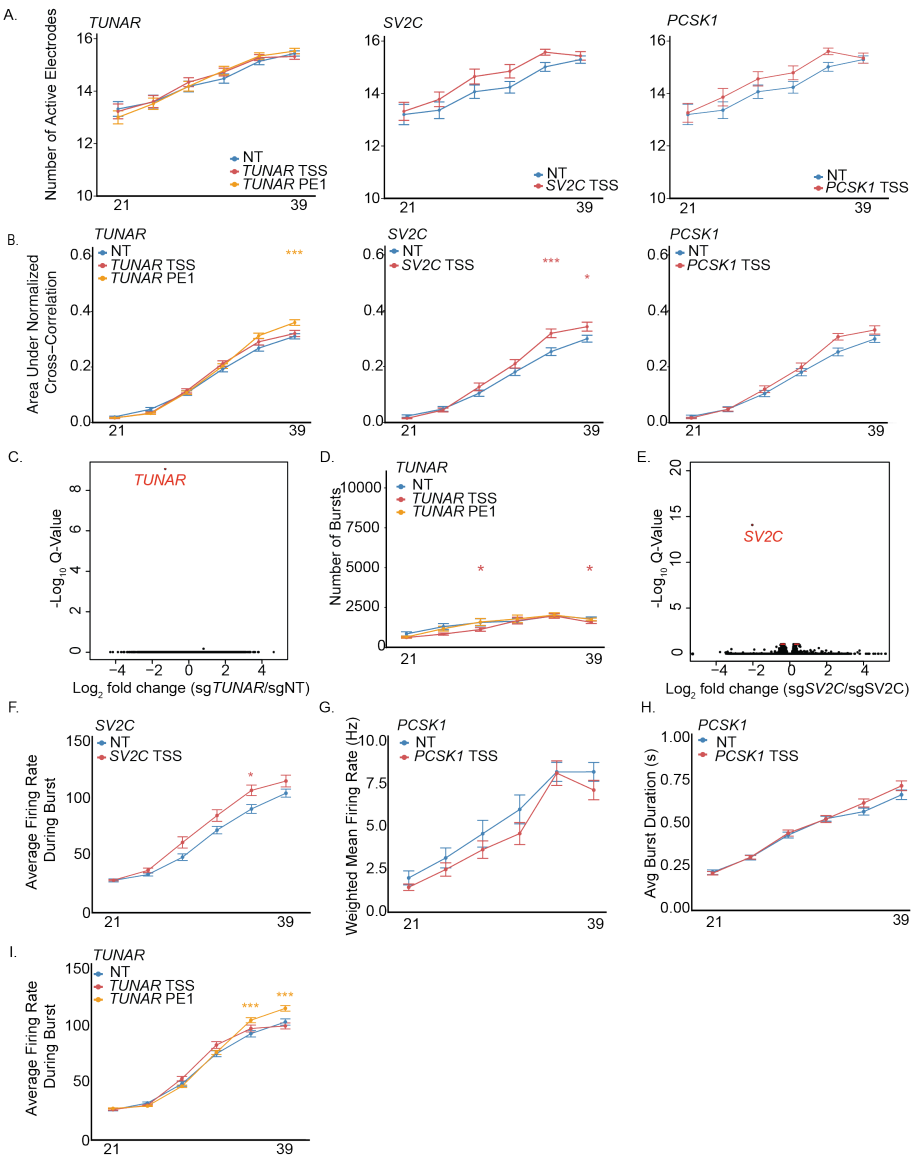


Figure S7: **Related to Figure 6.** a. Plot of number of active electrodes on MEA for NT, *TUNAR* TSS, *TUNAR* PE1, *SV2C* TSS, and *PCSK1* TSS CRISPRi lines recorded every 3/4 days from Day 21 to Day 39. Error bars are ± SEM. b. Plot of area under the normalized cross-correlation between electrodes on MEA for NT, *TUNAR* TSS, *TUNAR* PE1, *SV2C* TSS, and *PCSK1* TSS CRISPRi lines recorded every 3/4 days from Day 21 to Day 39. Error bars are ± SEM. c. Volcano plot of gene expression differences between *TUNAR* TSS and NT CRISPRi lines collected at Day 28. d. Plot of number of bursts on MEA for NT, *TUNAR* TSS, and *TUNAR* PE1 CRISPRi lines recorded every 3/4 days from Day 21 to Day 39. Error bars are ± SEM (* $p < 0.05$; Holm-Sidak corrected p-values from t-tests on estimated marginal means derived from mixed effects models). e. Volcano plot of gene expression differences between *SV2C* TSS and NT CRISPRi lines collected at Day 28. f. Plot of average firing rate during burst on MEA for NT and *SV2C* TSS CRISPRi lines recorded every 3/4 days from Day 21 to Day 39. Error bars are ± SEM (* $p < 0.05$; Holm-Sidak corrected p-values from t-tests on estimated marginal means derived from mixed effects models). g. Plot of weighted mean firing rate in *PCSK1* TSS, and NT CRISPRi lines recorded every 3/4 days from Day 21 to Day 39. Error bars are ± SEM. h. Plot of burst duration in *PCSK1* TSS, and NT CRISPRi lines recorded every 3/4 days from Day 21 to Day 39. Error bars are ± SEM. i. Plot of average firing rate during bursts on MEA for NT, *TUNAR* TSS, and *TUNAR* PE1 CRISPRi lines recorded every 3/4 days from Day 21 to Day 39. Error bars are ± SEM (*** $p < 0.001$; Holm-Sidak corrected p-values from t-tests on estimated marginal means derived from mixed effects models).

**Supplemental Tables**

Table S1: **Cell lines and oligos used in this study.**
Table S2: **RNA-seq analysis of human and chimpanzee diploid, autotetraploid, and allotetraploid piNs.**
Table S3: **ATAC-seq analysis of human and chimpanzee diploid, autotetraploid, and allotetraploid piNs.**
Table S4: **Analysis of CUT&Tag data (FOS and H3K27Ac)**
Table S5: **RNA-seq of piNs with *TUNAR* or *SV2C* CRISPRi**

### References

Malik, A. N., Vierbuchen, T., Hemberg, M., Rubin, A. A., Ling, E., Couch, C. H., Stroud, H., Spiegel, I., Farh, K. K.-H., Harmin, D. A., *et al.*, 2014. Genome-wide identification and characterization of functional neuronal activity-dependent enhancers. *Nature Neuroscience*, **17**(10):1330–1339.

Nehme, R., Zuccaro, E., Ghosh, S. D., Li, C., Sherwood, J. L., Pietilainen, O., Barrett, L. E., Limone, F., Worringer, K. A., Kommineni, S., *et al.*, 2018. Combining ngn2 programming with developmental patterning generates human excitatory neurons with nmdar-mediated synaptic transmission. *Cell Reports*, **23**(8):2509–2523.
